# Benchmarking ambient RNA removal across droplet and well-plate platforms reveals artificial count generation as a critical failure mode of scAR and CellClear

**DOI:** 10.64898/2026.04.08.717130

**Authors:** Lukas Schroeder, Susanne Gerber, Nicolas Ruffini

## Abstract

**Background:** Ambient RNA contamination is a pervasive artifact of single-cell and single-nucleus RNA sequencing (sxRNA-seq), yet no consensus exists on which computational removal tool performs best across experimental platforms.

**Results:** We present a systematic benchmark of six tools: CellBender, DecontX, SoupX, scCDC, scAR, and CellClear - evaluated across six human-mouse cell line mixing (hgmm) datasets (1k-20k cells) providing partial ground truth, two droplet-based complex tissue datasets (PBMC scRNA-seq; prefrontal cortex snRNA-seq), and a well-plate-based dataset (BD Rhapsody WBC). Using inter-species counts as partial ground truth, we quantify sensitivity, specificity, precision, and removal consistency per tool. We further apply a count-integrity criterion quantifying gene-cell positions where corrected values exceed raw counts. This reveals that scAR and CellClear do not merely denoise but fundamentally restructure count matrices: CellClear replaces >93% of counts with values derived from matrix factorization, while scAR generates spurious cell types absent from uncorrected data, including three spurious coarse cell types in the BD Rhapsody dataset and up to eight novel cell types in the prefrontal cortex. CellBender and SoupX exhibit reliable contamination removal with minimal count distortion. DecontX and scCDC are the only tools operable on non-droplet platforms without raw count matrix access. Runtime benchmarking at atlas scale (up to 172,000 nuclei) further demonstrates that CellClear fails to scale.

**Conclusions:** Count matrix integrity, not removal sensitivity alone, must be a primary criterion when selecting ambient RNA correction tools. We provide platform-specific recommendations and a decision framework to guide tool selection across experimental contexts.

## Background

### The ambient RNA artifact: origins and downstream consequences

In droplet-based sxRNA-seq, cell-free RNA present in the cell suspension is co-encapsulated with intact cells during droplet formation (Figure 1A). This ambient RNA pool arises primarily from cells lysed during tissue dissociation, and its composition reflects the transcriptional profile of the most abundant or fragile cell types in the sample (1,2). Because ambient RNA is captured by the same bead barcode as a cell’s own mRNA, it contaminates the resulting count matrix in a manner that is largely invisible without additional experimental or computational information. The consequences for downstream analysis are well-documented: ambient RNA reduces the specificity of marker genes, introduces spurious co-expression between transcriptionally distinct cell populations, confounds differential expression analysis when contamination levels differ between conditions or samples, and can produce apparent novel cell populations that reflect the ambient profile rather than genuine biology (3,4). Contamination is highly variable: Janssen et al. (3) observed 3-35% of total UMIs per cell across five mouse kidney replicates, with higher levels in snRNA-seq than matched scRNA-seq preparations, consistent with the greater mechanical stress of nuclei isolation and the shedding of ribosome-associated RNA from isolated nuclei into the loading buffer (5). Ambient contamination is not restricted to droplet-based platforms: in well-plate-based systems such as BD Rhapsody, cross-well contamination and extracellular RNA contribute ambient signal through distinct physical mechanisms, and Colino-Sanguino et al. (6) demonstrated explicitly that the source and character of ambient noise differ between plate-based and droplet-based platforms, constituting a distinction with direct implications for tool applicability.

**Figure 1:**
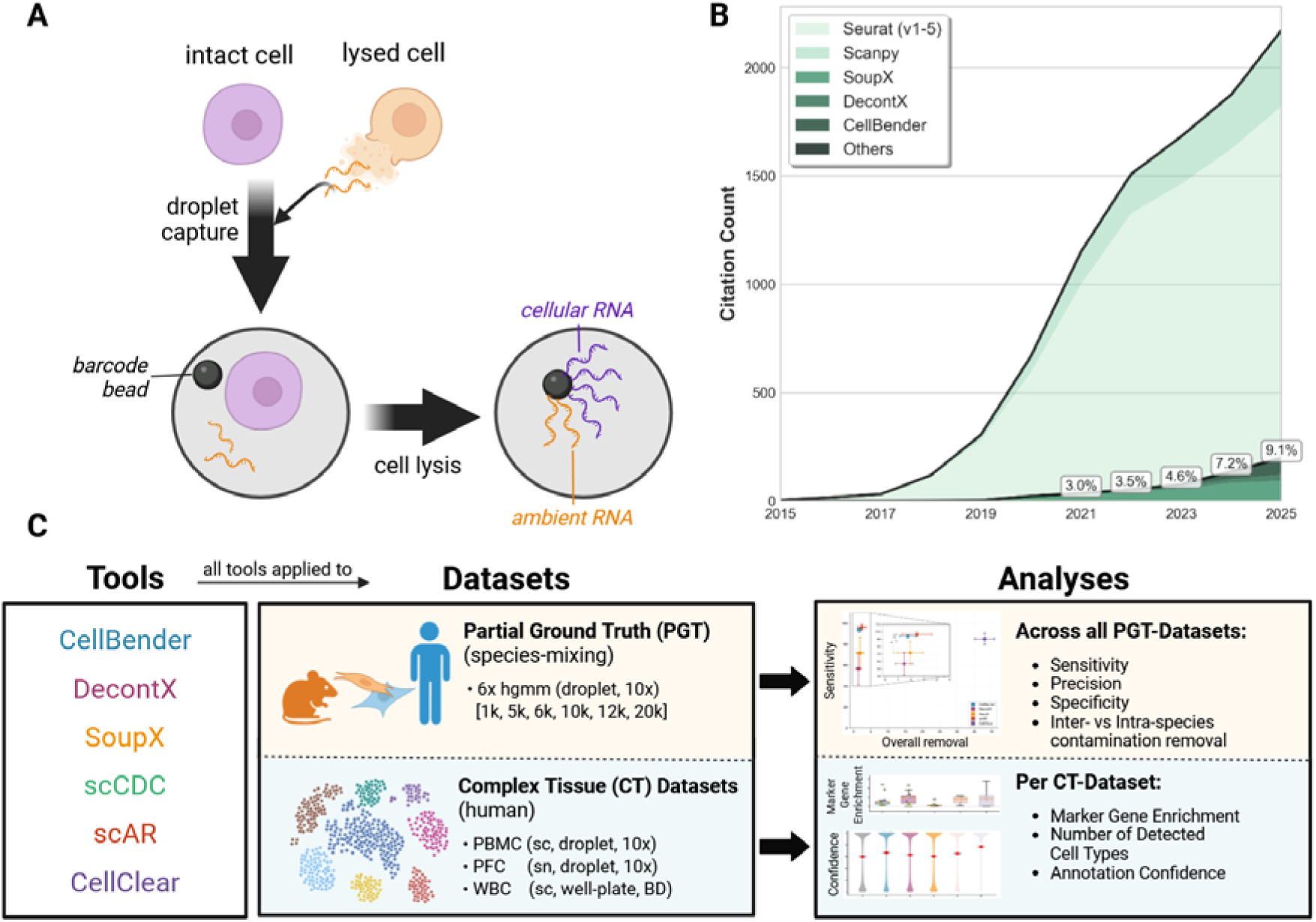
Ambient RNA contamination in sxRNA-seq and benchmarking study overview. **A)** Schematic of ambient RNA contamination (exemplified on scRNA seq data in droplet-based systems). During tissue dissociation, a proportion of cells lyse and release RNA into the suspension. Upon droplet capture, an intact cell is encapsulated together with ambient RNA from the surrounding solution. Following cell lysis and barcoding, both cellular RNA (purple) and ambient RNA (orange) are captured by the barcode bead, resulting in contaminated count matrices. **B)** Adoption of ambient RNA removal tools relative to overall scRNA-seq usage over time. Citation counts were retrieved via PubMed using the NCBI Entrez E-utilities API (Biopython v1.86). Papers citing Seurat (v1-5) or Scanpy and additionally reporting an associated GEO dataset entry were used as a proxy for empirical scRNA-seq studies (n=8,682). The fraction of these papers additionally citing SoupX, DecontX, CellBender or other (FastCAR / scCDC) ambient removal tools is shown per publication year. Percentages denote the annual adoption fraction per timepoint. **C)** Study design overview. All six tools (CellBender, DecontX, SoupX, scCDC, scAR, CellClear) were evaluated on two dataset types: species-mixing datasets providing partial ground truth (PGT; 6 hgmm datasets, 1k-20k cells) and complex tissue (CT) datasets (PBMC, PFC, WBC). Performance was assessed across multiple metrics including sensitivity, precision, specificity for partial ground truth datasets and metrices such as marker gene enrichment, confidence of cell type identification and number of confidently identified cell types for complex tissue datasets. Created in BioRender. Ruffini, N. (2026) https://BioRender.com/4xtbbmm.

Ambient RNA removal has emerged as a near-routine preprocessing step within many of these workflows: integration into frameworks such as nf-core (7) has lowered the barrier to applying probabilistic background correction, and adoption of dedicated tools has risen steadily alongside overall sxRNA-seq usage (Figure 1B). Yet this routinization has outpaced independent evaluation. The last peer-reviewed benchmark of ambient removal tools (3) pre-dates the publication of several newer tools: scAR (8), scCDC (9), CellClear (10), and FastCAR (11), the latter developed specifically for differential gene expression optimization rather than global ambient removal and was restricted to three tools on a single experimental platform. As tools with fundamentally different algorithmic architectures enter routine use with varying levels of scrutiny, an updated and broader benchmark is warranted.

### A diverse computational landscape for ambient RNA removal

Six tools have achieved sufficient adoption to merit systematic evaluation here (Figure 1C, Table 1). CellBender (2) employs a variational autoencoder (VAE) to jointly model ambient signal and cell-specific expression from both empty and non-empty droplets, requiring the raw count matrix and benefiting substantially from GPU acceleration. DecontX (4) models each cell as a Bayesian mixture of native and contaminating counts using cluster assignments; it operates without empty droplet data and without requiring the raw unfiltered count matrix, rendering it applicable to non-droplet platforms and to pre-processed count matrices as they are frequently encountered in public data deposits. SoupX (1) estimates a global contamination fraction via maximum likelihood using empty droplet profiles and applies proportional subtraction across all genes, requiring the raw unfiltered count matrix. scCDC (9) similarly operates on pre-filtered count matrices without requiring empty droplets, but applies gene-selective correction only to detected contamination-causing genes rather than performing global ambient removal. It was developed and validated exclusively on droplet-based data, however, as it requires neither empty droplets nor a raw unfiltered count matrix, its authors claim general applicability to publicly deposited scRNA-seq and snRNA-seq datasets regardless of platform. While this paradigm is methodologically distinct from global ambient removal, we include scCDC as a reference comparator given its adoption in the field, with the caveat that its gene-selective design means sensitivity and precision metrics reflect a narrower correction target than the other tools evaluated. Similar to CellBender, scAR (8) uses a VAE architecture under the ambient signal hypothesis, modelling per-cell noise ratios and native expression frequencies as latent variables; it was designed as a technology-agnostic tool intended for scRNA-seq, CITE-seq, and CRISPR screen data alike. CellClear (10) also performs per-gene rather than global correction, but via a structurally distinct mechanism: it applies non-negative matrix factorization (NMF) to foreground and background count matrices to identify genes whose foreground-to-background distance is anomalous, and uses AUROC-based thresholding to determine which genes undergo correction. However, unlike scCDC’s targeted subtraction, CellClear’s NMF decomposition reconstructs the entire count matrix, with the consequence that corrected values are derived from factorized rather than original counts across all genes. One additional tool was considered but excluded: FastCAR (11) applies gene-level threshold subtraction optimized specifically for differential gene expression (DGE) analysis rather than global ambient removal, making it incompatible with the sensitivity and specificity metrics derived from species-mixing ground truth used here.

**Table 1:**
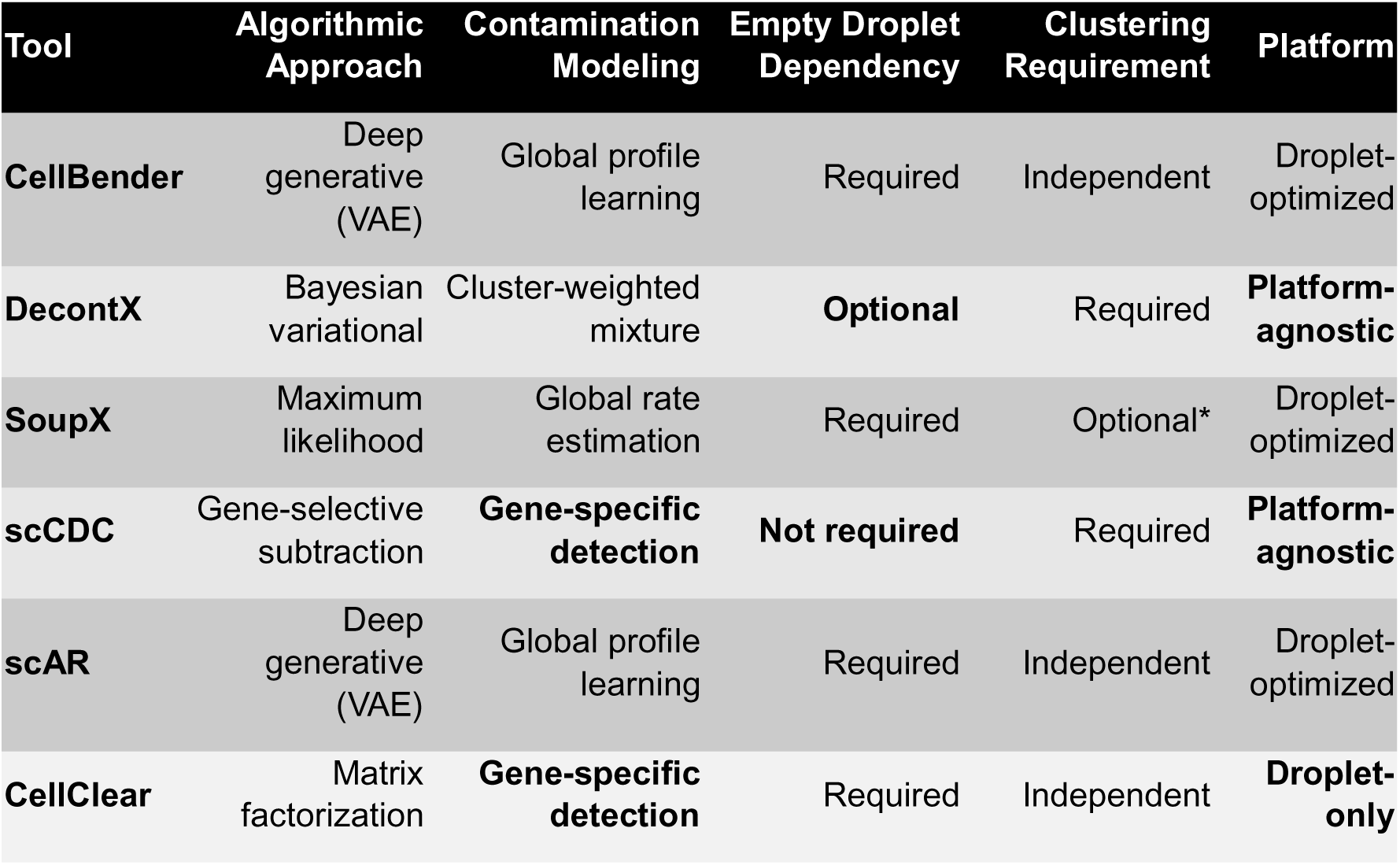
Overview of evaluated ambient RNA correction tools. Key algorithmic and practical properties are summarized for each method, including the underlying modeling approach, how the ambient RNA profile is estimated, whether empty droplets are required as input, whether clustering information is needed, and the intended sequencing platform. *Clustering information used for improved ambient profile estimation but not algorithmically required.

| Tool | Algorithmic Approach | Contamination Modeling | Empty Droplet Dependency | Clustering Requirement | Platform |
| --- | --- | --- | --- | --- | --- |
| **CellBender** | Deep generative (VAE) | Global profile learning | Required | Independent | Droplet-optimized |
| **DecontX** | Bayesian variational | Cluster-weighted mixture | **Optional** | Required | **Platform-agnostic** |
| **SoupX** | Maximum likelihood | Global rate estimation | Required | Optional* | Droplet-optimized |
| **scCDC** | Gene-selective subtraction | **Gene-specific detection** | **Not required** | Required | **Platform-agnostic** |
| **scAR** | Deep generative (VAE) | Global profile learning | Required | Independent | Droplet-optimized |
| **CellClear** | Matrix factorization | **Gene-specific detection** | Required | Independent | **Droplet-only** |

### Prior benchmarks and their limitations

The most rigorous prior peer-reviewed benchmark, Janssen et al. (3), evaluated CellBender, DecontX, and SoupX using mouse kidney datasets in which cells from two mouse subspecies could be discriminated via known homozygous SNPs, providing ground truth in a genuinely complex, multi-cell-type setting. They found CellBender to yield the most precise ambient estimates and the greatest improvement in marker gene detection, while clustering and cell classification were relatively robust to background noise regardless of the tool applied. This work established an important baseline but was restricted to three tools, one tissue type, and droplet-based platforms exclusively. The recent bioRxiv preprint by Cargnelli et al. (12) subsequently expanded the evaluation to seven tools, adding scAR, scCDC, FastCAR, and CellClear, and included simulated datasets with known contamination levels, genotype-mixing experiments, and negative control datasets using SmartSeq2-based data. Cargnelli et al. found no single method superior across all metrics and noted that scAR performs poorly on negative controls and leads to worse batch integration, while CellClear shows limited overall ambient removal relative to the extent of count alteration it produces. However, neither benchmark included a well-plate-based platform, and neither explicitly quantified artificial count inflation, i.e. the systematic generation of non-zero corrected values at positions with zero raw counts, as a distinct failure mode. The dataset panel in both prior benchmarks also did not span the range of contamination levels and cell numbers needed to characterize performance variability comprehensively.

### This study

Here, we present a systematic benchmark of six ambient RNA removal tools: CellBender, DecontX, SoupX, scCDC, scAR, and CellClear, across a broader and more diverse dataset panel than any prior benchmark (Figure 1C). All tools were evaluated on six publicly available human-mouse cell line mixing (hgmm) datasets from 10x Genomics spanning a tenfold range of cell numbers (1k-20k cells), providing partial quantitative ground truth for sensitivity, specificity, precision, and removal consistency. We additionally evaluated all tools on three complex tissue datasets, two of which being droplet-based (PBMC scRNA-seq & prefrontal cortex (PFC) snRNA-seq dataset) and one being a well-plate-based BD Rhapsody white blood cell (WBC) dataset, extending benchmark scope to a non-droplet platform not addressed in prior studies. We further investigate artificial count increment by quantifying the fraction of final corrected count values that exceed the corresponding raw counts on a per-cell, per-gene basis, revealing the extent to which corrected matrices contain artificial signal not present in the original data and assess the downstream consequences of this count restructuring through automated cell type annotation across all three complex tissue datasets. Computational scalability was evaluated across datasets ranging from 18,000 to 172,000 cells and nuclei. Together, these analyses establish count matrix integrity, rather than removal sensitivity alone, as the decisive criterion for ambient correction tool selection, and yield platform-specific recommendations and a decision framework for practitioners.

## Results

Results are organized in four analytical layers. We first quantify ambient RNA removal performance using species-mixing ground truth across six hgmm datasets (Figure 2). We then assess downstream effects on cell type annotation quality in three complex tissue datasets (Figure 3). Two tools, namely scAR and CellClear, are excluded from statistical inference in Figure 2 and 3 on the basis of systematic count inflation, the characterization of which constitutes the third analytical layer (Figure 4). Finally, computational scalability and practical tool selection are summarized in Figure 5. The count inflation analyses in Figure 4 are presented after partial ground-truth and the annotation results because the annotation findings motivate and contextualize the inflation analyses, but readers may wish to consult Figure 4 before Figure 2 and 3.

**Figure 2:**
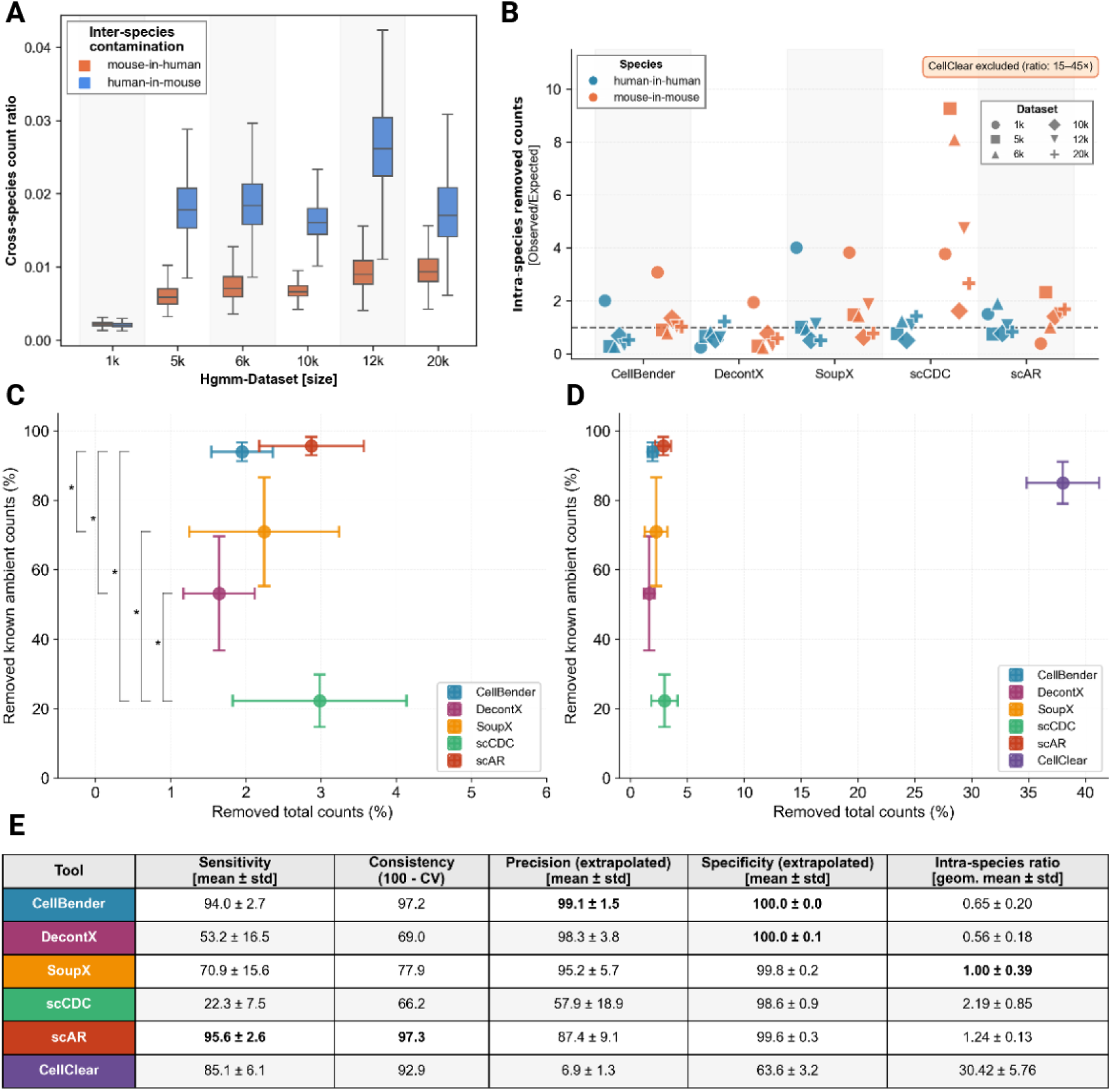
Ambient RNA removal performance across six hgmm species-mixing datasets. **A)** Baseline cross-species contamination (inter-species count ratio) across all six hgmm datasets before correction, stratified by direction (mouse-in-human, human-in-mouse). The 1k dataset exhibits markedly lower ambient contamination than larger datasets. **B)** Intra-species removal ratio (observed/expected) per tool and dataset for CellBender, DecontX, SoupX, scCDC and scAR (CellClear excluded; observed ratio 15-45× above expected). The dashed reference line at 1.0 indicates proportional removal relative to the extrapolated true intra-species ambient load. **C)** Tradeoff between total count removal and sensitivity (removed known ambient counts), summarized as mean ± SD across five datasets (1k excluded), shown for all tools except CellClear. Asterisks indicate significant pairwise differences in sensitivity (p < 0.05, Wilcoxon signed-rank, Supp. Table 1,2). **D)** Same data as C including CellClear, shown on full x-axis scale. CellClear’s extreme total count removal (∼35-40%) compresses all other tools into the leftmost portion of the axis. **E)** Summary metrics table aggregated across five datasets (1k excluded). Sensitivity: fraction of known ambient counts removed. Consistency: 100 minus the coefficient of variation of sensitivity. Precision and specificity are extrapolated accounting for undetectable intra-species contamination. Intra-species ratio: geometric mean of observed-to-expected removal for human-in-human and mouse-in-mouse counts. Bold values indicate best performance per metric.

**Figure 3.**
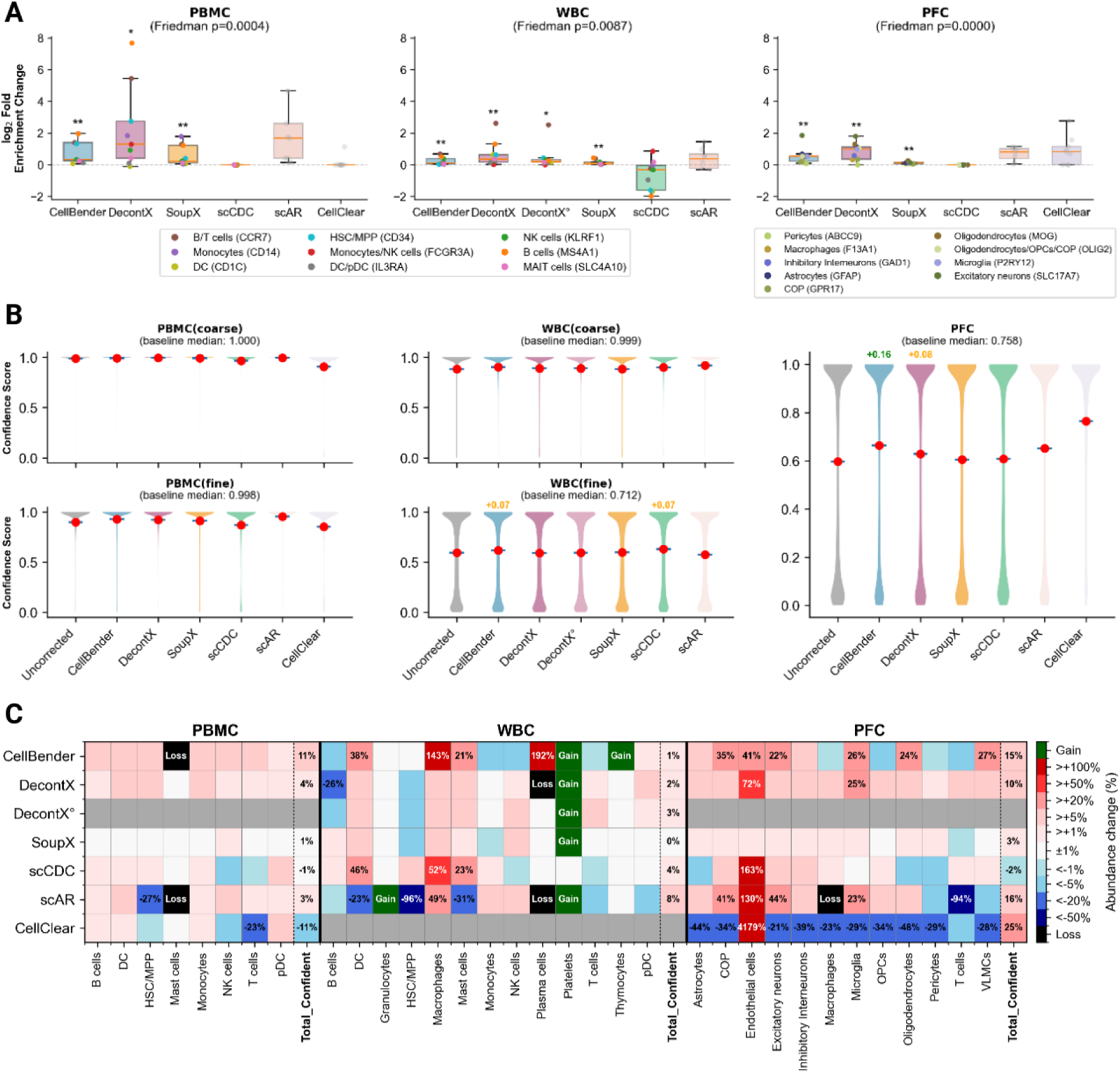
Effects of ambient RNA correction on cell type annotation quality across three complex tissue datasets. **A)** Marker gene log□-fold enrichment ratio change after ambient RNA correction for PBMC, WBC, and PFC datasets. Each data point represents one marker gene’s log_2_-fold enrichment post-vs pre-correction regarding its cell counts across cells of the expected cell vs its counts in all other cells. Boxplots show the distribution across markers for all analyzed tools. Significance annotations above each tool indicate one-sided Wilcoxon signed-rank tests against zero (H□: median > 0), Benjamini-Hochberg-corrected across tools within each dataset; Friedman test p-values are shown in panel titles and Supp. Table 3. scAR and CellClear were excluded from statistical testing due to systematic count inflation invalidating ratio-based comparisons. (see Results) **B)** Distribution of CellTypist confidence scores per cell with and without ambient RNA correction. Violin plots are shown for coarse and fine annotation models where applicable. Red dots indicate mean confidence scores. Annotated values indicate median differences exceeding 0.05 relative to uncorrected baseline. scAR and CellClear are shown in muted colors as they were excluded from statistical inference due to count inflation artifacts. **C)** Heatmap of relative cell type abundance changes after ambient RNA correction compared to uncorrected counts. Gains (newly detected cell types) and losses (complete disappearance) are annotated explicitly; percentage changes > 20% or for the Total_Confident summary column are labeled. Grey cells indicate tool-dataset combinations that were not applicable. Bold column labels denote the Total_Confident summary column.

**Figure 4:**
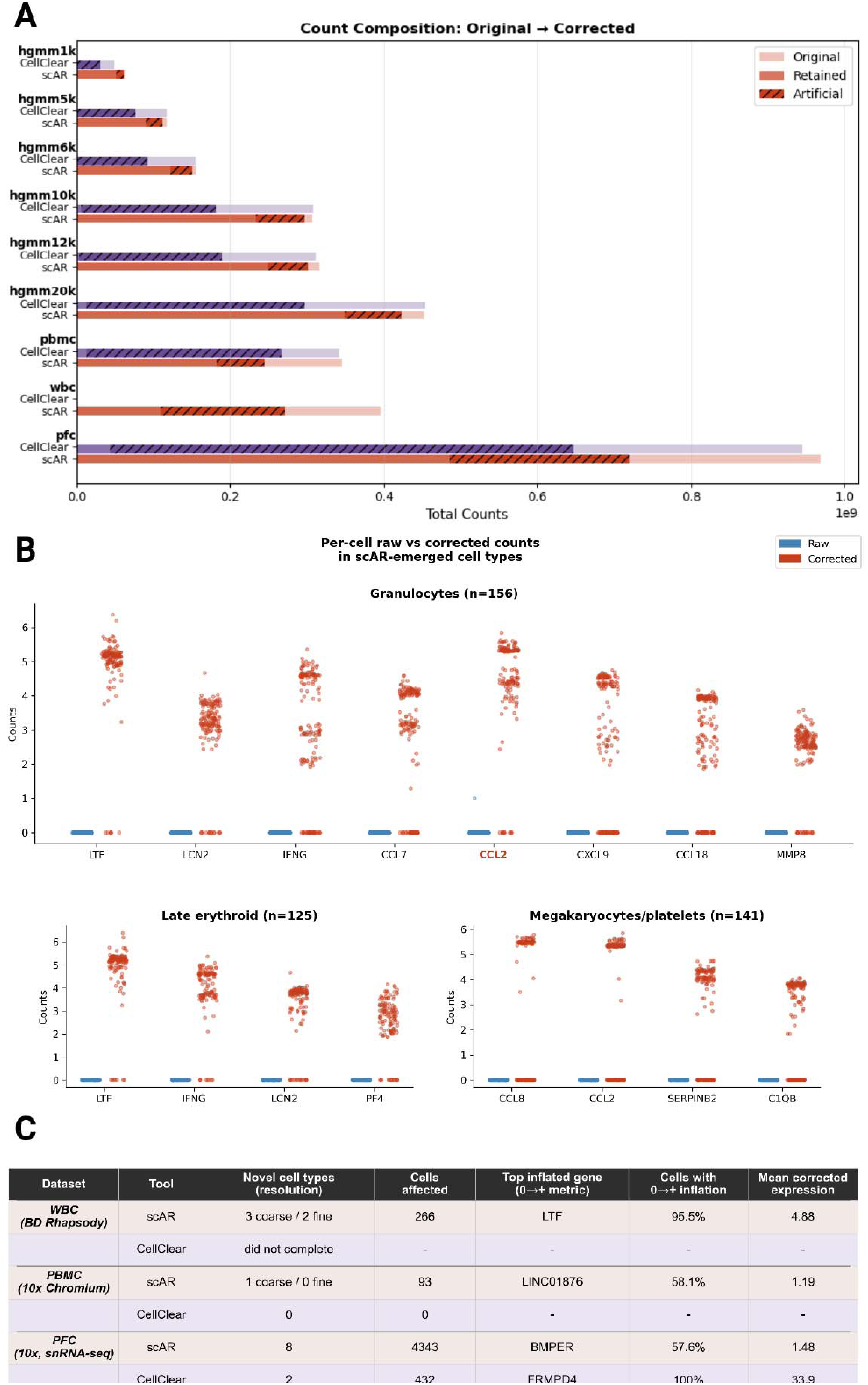
scAR and CellClear introduce artificial counts and generate spurious cell types. **A)** Count composition before and after ambient RNA correction with scAR and CellClear across all benchmarked datasets. For each dataset, bars show original total counts (light), counts retained after correction (solid), and counts artificially inflated above original values (hatched). Datasets are grouped by biological context (PFC, WBC, PBMC) and six human-mouse cell line mixture datasets (hgmm) of increasing cell numbers. **B)** Per-cell uncorrected (blue) versus scAR-corrected (red) counts for the top inflated genes in three cell types absent from the uncorrected WBC (BD Rhapsody) dataset but emerged after scAR correction. Horizontal lines indicate per-gene means. Genes highlighted in red are canonical markers for the respective cell type. All displayed raw counts are near zero, confirming artificial inflation rather than denoising. **C)** Summary of novel cell types emerging after correction with scAR or CellClear across biological datasets. Reported are the number of cell types gained at coarse and fine annotation resolution, total cells affected, the top gene by the zero-to-positive inflation metric (highest proportion of cells transitioning from zero raw counts to non-zero corrected counts), the fraction of cells with such zero-to-positive inflation, and the mean corrected expression of that gene after artificial inflation.

**Figure 5:**
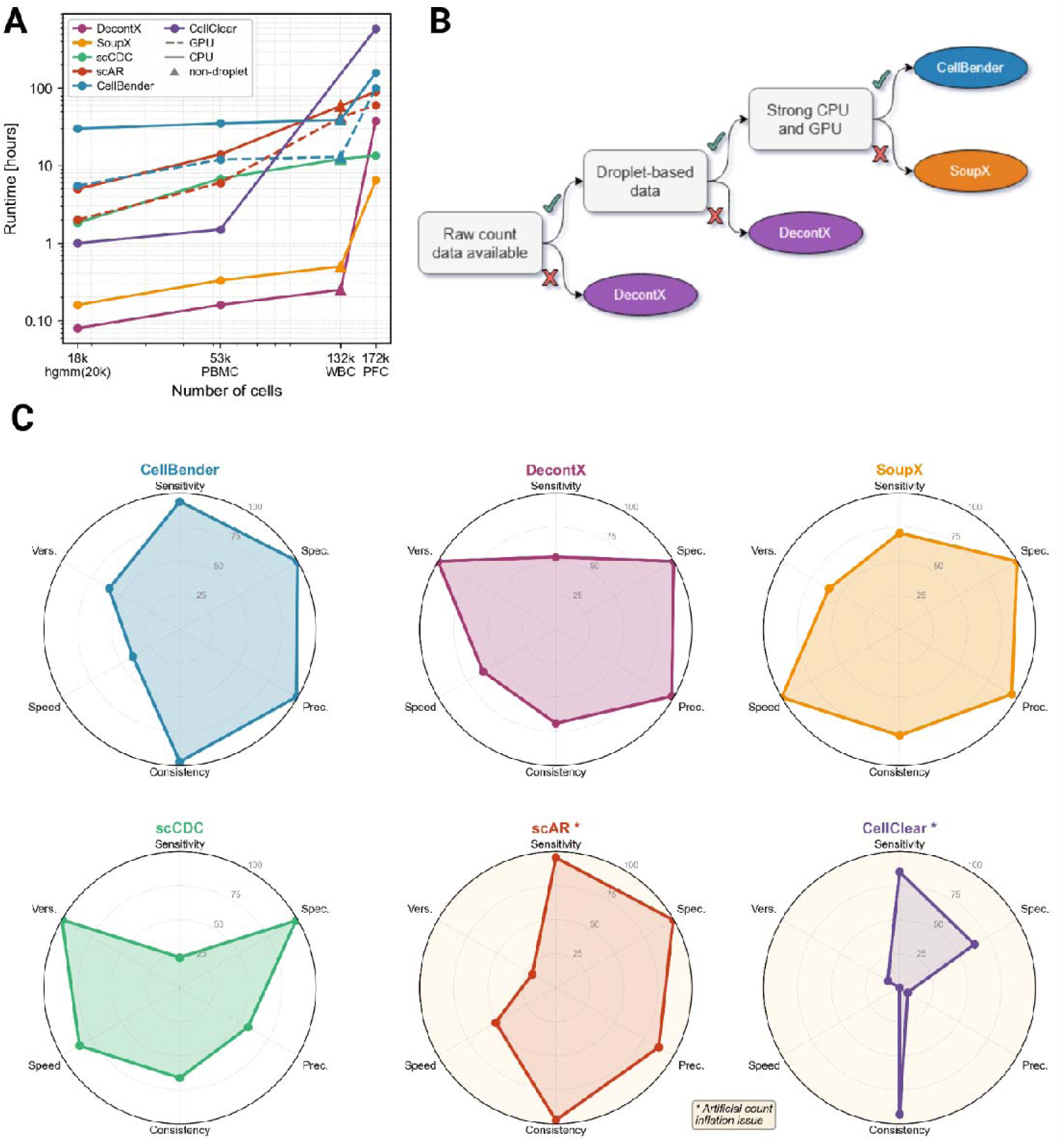
Scalability, compatibility and overall performance of ambient RNA correction tools. **A)** Wall-clock runtimes across datasets of increasing cell number (18,000-172,000). Times include file I/O, correction, and output writing. Machine learning-based tools (scAR, CellBender) are shown for both CPU and GPU execution. CellClear lacks a datapoint for the WBC dataset as it failed for the well-plate based platform. The WBC triangle marker indicates non-droplet (well plate-based) data. Y-axis is log-scaled. **B)** Decision tree summarizing recommended tool selection based on platform type, raw count data availability, and computational resources. scAR and CellClear are excluded from recommendations due to artificial count inflation demonstrated in Figure 4. **C)** Radar plots summarizing tool performance across six dimensions. Sensitivity, Specificity, Precision, and Consistency are derived from hgmm benchmarking (Figure 2E). Speed is computed as the log-scale min-max normalized inverse of maximum observed runtime, anchored at SoupX (6.5 h = 100) and CellClear (582 h = 0). Versatility reflects operational scope: DecontX and scCDC both received the maximum score (100) as they run without raw count matrices and are applicable to non-droplet platforms. CellBender and SoupX received 60 reflecting droplet-platform dependency and raw data requirements; scAR and CellClear received 20 and 10 respectively, additionally penalized for artificial count inflation that critically limits the utility of their output across all downstream analyses. Asterisks denote tools not recommended for routine use (see Figure 4).

Baseline inter-species contamination varied substantially across hgmm datasets (Figure 2A). The 1k dataset exhibited markedly lower ambient counts than all larger datasets. This pattern is consistent with fewer cell lysis events in smaller capture volumes and lower total cell stress. As this outlier behavior propagates into the intra-species extrapolation (Figure 2B), where the 1k dataset produces anomalously high ratios across multiple tools, it is the primary reason for its exclusion from all quantitative aggregations. For non-1k datasets, inter-species contamination increased with dataset size, with the human-in-mouse direction consistently exceeding mouse-in-human, likely reflecting the higher fragility of HEK293T cells relative to NIH3T3.

Intra-species removal ratios varied considerably across tools (Figure 2B). DecontX was the most conservative, with ratios consistently below 1.0 across all non-1k datasets and both species, indicating systematic under-removal relative to the expected intra-species ambient load. SoupX clustered near 1.0 for non-1k datasets (geometric mean 1.00 ± 0.39), indicating roughly proportional removal, though with notable variance across datasets. CellBender showed a similar central tendency to SoupX among non-1k datasets, with most points near or slightly below 1.0. scAR showed elevated ratios among non-1k datasets (geometric mean 1.24 ± 0.13), with multiple larger datasets exceeding 1.0 in both species directions, suggesting removal beyond what the ambient load alone accounts for, foreshadowing a pattern addressed further in Figure 4. scCDC exhibited the highest overall intra-species ratio (2.19 ± 0.85) but with a pronounced species asymmetry: mouse intra-species removal was strongly disproportionate to the expected ambient load, while human intra-species removal remained near 1.0. This asymmetry indicates that scCDC’s gene selection is heavily biased toward mouse-origin genes in this dataset context, consistent with selective targeting of the most readily detectable contamination signal rather than proportional ambient profile removal.

CellBender and scAR achieved the highest mean sensitivity across non-1k datasets (94.0 ± 2.7% and 95.6 ± 2.6% respectively; Figure 2C-E). scAR showed superior consistency (97.3 vs 97.2 for CellBender); this apparent advantage must however be interpreted alongside its count inflation behavior documented in Figure 4. SoupX and DecontX were substantially more conservative, removing 70.9 ± 15.6% and 53.2 ± 16.5% of known ambient counts respectively, with DecontX exhibiting lower consistency (69.0) driven by sensitivity variance across datasets. scCDC achieved the lowest mean sensitivity of all tools (22.3 ± 7.5%) and low consistency (66.2), in line with its design objective of targeting only high-confidence contamination-causing genes rather than performing global ambient removal.

CellBender and DecontX achieved the highest extrapolated specificity (100%), while SoupX and scAR were comparable (99.8% and 99.6%). scCDC’s extrapolated precision of 57.9% reflects the species asymmetry in its gene selection, with a substantial fraction of removed counts deriving from endogenous rather than ambient signal. CellClear achieved 85.1% mean sensitivity but at the cost of removing 35-40% of total counts; an order of magnitude more than any other tool. Its extrapolated precision of 6.9% and specificity of 63.6% confirm that the vast majority of counts removed were not ambient RNA.

Taken together, CellBender offers the best combination of sensitivity, precision, and specificity among tools without count inflation artifacts. SoupX performs near the expected intra-species removal ratio with high specificity, at the cost of lower sensitivity. DecontX is the most conservative tool, systematically under-removing relative to the extrapolated ambient load. scCDC’s low sensitivity and species-asymmetric removal pattern are consistent with its gene-selective design but preclude its use as a substitute for global ambient correction. CellClear’s precision and specificity render it unsuitable for reliable ambient correction despite its high apparent sensitivity.

Ambient RNA correction significantly improved marker gene enrichment ratios across all three datasets (Friedman test: PBMC χ² = 18.47, p = 0.0004; WBC χ² = 13.60, p = 0.0087; PFC χ² = 23.13, p < 0.0001). Among the four tools included in statistical testing, DecontX consistently achieved the highest median enrichment changes (PBMC: +1.30 log□FC, WBC: +0.35, PFC: +0.98), followed by CellBender, with SoupX showing the most conservative but uniformly positive improvement. scCDC produced near or below zero median enrichment changes across all three datasets (PBMC & PFC: +0, WBC: −0.319), consistent with its gene-selective design targeting only high-confidence contamination-causing genes rather than performing global ambient removal. DecontX run with background estimation (DecontX) and without (DecontX°: using only the filtered count matrix) showed comparable enrichment patterns, with no significant pairwise difference in the WBC dataset (pairwise Wilcoxon, p = 0.074, Supp. Table 4). This suggests that DecontX’s correction efficacy is not critically dependent on access to the raw count matrix in this dataset context. Post-hoc comparisons showed DecontX significantly outperformed SoupX in PBMC and PFC (p = 0.020 and p = 0.008), and CellBender outperformed SoupX in PFC (p = 0.004); CellBender outperformed DecontX in PFC (p = 0.039) but not in PBMC or WBC. All three global-correction tools significantly outperformed scCDC in PBMC and PFC (all p ≤ 0.012); in WBC, only DecontX and DecontX° differed significantly from scCDC (p = 0.027 and p = 0.039), while CellBender and SoupX did not reach significance (p = 0.055 and p = 0.074; Supplementary Table 4). scAR and CellClear were excluded from statistical testing, as scAR produces nominally elevated enrichment ratios that are uninterpretable in the context of its count inflation behavior (as documented in Figure 4). CellClear produced near-zero enrichment changes in PBMC and WBC, consistent with its limited effective correction scope despite extreme total count alteration (Figure 4).

The most compelling evidence for the practical utility of ambient correction in complex tissue data came from cell type composition analysis. Platelets were undetectable in uncorrected WBC data but were confidently recovered by every correction tool evaluated - except for scCDC - establishing that their absence prior to correction reflects ambient RNA suppression of canonical platelet markers rather than true biological absence or any tool-specific artifact (Figure 3C). CellBender additionally enabled confident detection of Thymocytes in WBC, a rarer population not recovered by any other tool.

CellTypist (13) confidence scores revealed ceiling effects in PBMC and WBC coarse annotation, where baseline medians of 1.000 and 0.999 left virtually no room for improvement. All tools showed statistically significant but negligible effects in these settings (all Cliff’s δ < 0.147). Under the WBC fine annotation model, the lower baseline median (0.707) allowed CellBender to produce a median increase of +0.069; scCDC produced a comparable increase of +0.067, while DecontX° and SoupX showed no substantial (> +0.05) median improvement. Notably, scCDC produced statistically significant but negligible *decreases* in annotation confidence relative to uncorrected in both PBMC annotation models (Cliff’s δ = −0.068 and −0.065, coarse and fine respectively; Supp. Table 5), and DecontX showed a marginal decrease under WBC fine annotation (δ = −0.008), suggesting that in the absence of meaningful ambient removal, correction can very slightly disorganize count distributions in ways that fractionally reduce classifier confidence.

The largest gains were observed in PFC, where higher ambient burden in snRNA-seq combined with a more granular cortical CellTypist model yielded a baseline median of 0.758. CellBender and DecontX produced median increases of +0.163 and +0.083 respectively, differences that are practically meaningful at this confidence scale (Figure 3B).

In PFC, the number of confidently annotated cells increased broadly across tools, with DecontX and CellBender showing the most consistent gains without implausible compositional shifts and SoupX again being the most conservative. scCDC produced largely unremarkable compositional shifts in PBMC and WBC but showed a 163% increase in confidently annotated Endothelial cells in PFC alongside near-zero or negative changes in most other cell types. This pattern is consistent with scCDC’s gene-selective design targeting a small number of highly expressed contamination-causing genes: in snRNA-seq of brain tissue, vascular-derived RNA is a known constituent of the ambient pool due to mechanical disruption of vasculature during nuclei isolation, and selective correction of such highly expressed endothelial markers would produce exactly the focal cell type abundance shift observed here without meaningful global ambient removal. scAR produced unexpected cell type gains and losses across datasets, including complete disappearance of Macrophages in PFC (n = 141 in uncorrected data) and Plasma cells in WBC - the mechanistic basis of which is examined in Figure 4. CellClear produced the most extreme compositional restructuring: in PFC, nearly every cell type contracted by 25-50% while Endothelial cells inflated 4,179-fold, showing a pattern incompatible with genuine denoising and addressed in Figure 4. Fine-resolution annotation results for PBMC and WBC, analogous to the fine-model results in Figure 3C, are shown in Supplementary Figure 1.

scAR and CellClear systematically altered the native count distribution beyond ambient RNA removal (Figure 4A). Across all datasets, CellClear removed >95% of original counts before reintroducing synthetic signal, resulting in corrected matrices with >93% artificial counts. scAR removed 18-72% of original counts while adding 17-60% artificial counts, with the largest effect in the WBC dataset.

This inflation had direct consequences for cell type annotation. In the WBC (BD Rhapsody) dataset, scAR correction produced three novel coarse cell types: Granulocytes (n=156), Megakaryocytes/platelets (n=69), and Early MK cells (n=41), none of which were detectable in uncorrected data. Inspection of per-cell counts confirmed that defining marker genes for these types had near-zero raw expression but were inflated to mean corrected values of 3-5 counts per cell (Figure 4B; Table in C). For example, LTF, a canonical granulocyte and erythroid marker, showed zero raw counts in >95% of affected cells yet reached a mean corrected expression of 4.88. In the PFC (snRNA-seq) dataset, scAR generated 8 novel fine cell types (n=4,343 cells total), while CellClear produced 2 (n=432), whose gene counts increment is further visualized in Supplementary Figure 2. The latter, CellClear, exhibited extreme inflation; FRMPD4 was inflated from zero to non-zero in 100% of cells in one emerged type, with a mean corrected expression of 33.9 (Figure 4C). In the PBMC dataset, scAR produced one spurious coarse cell type (HSC/MPP, n=93), while CellClear generated no novel types. The count inflation documented here also accounts retrospectively for scAR’s elevated intra-species removal ratios in Figure 2B: when scAR-corrected values are clipped per cell and gene to not exceed corresponding raw counts prior to metric calculation, ratios return to the range of CellBender and SoupX (Supplementary Figure 3), confirming that the Figure 2B signal reflects artificial count addition rather than genuine over-removal of endogenous signal. These results demonstrate that scAR and CellClear do not merely denoise but fundamentally restructure the count matrix, introducing biologically unsubstantiated cell identities.

Runtime scaling differed by orders of magnitude across tools (Figure 5A). SoupX was the fastest across all dataset sizes (maximum 6.5h). DecontX was faster on smaller datasets but showed disproportionate runtime increase on the 172,000-nucleus PFC dataset, likely reflecting memory pressure rather than algorithmic scaling behavior. scCDC occupied a middle tier, reaching 13.5h on the PFC dataset, with notably flat scaling between the 132,000-cell WBC dataset (12.2h) and PFC (13.5h), suggesting its gene-selective correction paradigm does not scale quadratically with cell number. CellBender scaled stably at moderate sizes but required 100h (GPU) and 158h (CPU) on the PFC dataset. scAR exhibited apparent quadratic scaling behavior, reaching 60.5h (GPU) and 90h (CPU) at maximum. CellClear completed in under 2h on smaller inputs but required 582h on PFC, confirming that it does not scale to atlas-size data in its current implementation. GPU acceleration reduced runtimes by approximately 45% for scAR and 60% for CellBender; as these measurements were obtained on a consumer-grade GPU (NVIDIA GTX 1050 Ti, 4 GB VRAM), absolute runtimes for both tools are likely overestimates relative to research-grade hardware, though relative tool comparisons remain valid.

Taken together, the hgmm benchmarking, annotation, count inflation, and scalability analyses identify three tools appropriate for routine use, with distinct operational niches (Figure 5B, 5C). For droplet-based data with GPU access, CellBender is the recommended primary choice given its superior sensitivity and precision. SoupX is the preferred lightweight alternative, offering robust removal, the fastest runtimes, and minimal count distortion, at the cost of lower sensitivity. DecontX and scCDC are the only tools in this panel applicable to non-droplet platforms and to datasets lacking raw unfiltered count matrices and are accordingly the sole appropriate option for the BD Rhapsody WBC dataset and for any pre-processed public dataset deposited without the raw count matrix. However, in terms of sensitivity and precision, DecontX outperforms scCDC. scAR and CellClear are excluded from recommendations on the basis of systematic artificial count introduction documented in Figure 4, representing a disqualifying characteristic independent of their contamination removal metrics.

Across all analytical layers, three properties emerge as decisive criteria for ambient correction tool selection: removal sensitivity, as quantified by hgmm ground truth; count matrix integrity, as assessed by the inflation analysis; and operational scope, as determined by platform and data availability constraints. These criteria produce a consistent and non-redundant ranking across tools, and their convergence with two independent benchmarks employing different ground truth strategies (3,12) strengthens confidence in the recommendations. The following Discussion contextualizes these findings in terms of the algorithmic origins of count inflation, the implications of the well-plate findings, and the broader question of when ambient correction is warranted.

## Discussion

### Count matrix integrity as the decisive benchmark criterion

A central finding of this benchmark is that removal sensitivity, the most commonly reported metric in ambient correction tool evaluations, is insufficient to discriminate safe from problematic tools. scAR and CellClear each demonstrate competitive or high apparent sensitivity in the hgmm benchmarks in the sense that ambient counts are removed, yet both systematically increase gene counts in the corrected matrix above their original raw values (Figure 4A), which is biologically and logically incompatible with the concept of ambient RNA removal. CellClear removes more than 95% of original counts before reintroducing values derived from NMF decomposition, resulting in corrected matrices in which more than 93% of counts are synthetic. scAR removes 18-72% of original counts across datasets while simultaneously adding 17-60% artificial counts via its VAE decoder, with the largest perturbation in the well-plate WBC dataset. The downstream consequences are not subtle. In the BD Rhapsody WBC dataset, scAR produced three coarse cell types entirely absent from uncorrected data, namely granulocytes (n=156), megakaryocytes/platelets (n=69), and early MK cells (n=41), whose defining marker genes, including LTF, had zero raw counts in more than 95% of affected cells yet reached mean corrected values of 3-5 counts per cell (Figure 4B, 4C). In the PFC snRNA-seq dataset, scAR generated eight novel fine-grained cell types totaling 4,343 cells, while CellClear produced two spurious types comprising 432 cells; in the PBMC dataset, scAR produced one spurious coarse cell type (HSC/MPP, n=93). This pattern of ambient contamination driving spurious cell type annotation in brain snRNA-seq is consistent with Caglayan et al. (14), who showed that neuronal ambient RNA pervasively contaminates glial nuclei and that a previously published immature oligodendrocyte population was entirely an ambient RNA artifact. The severity of the scAR artifact in the WBC dataset is consistent with its generative model encountering input statistics outside its training distribution, a concern directly supported by Colino-Sanguino et al. (6), who demonstrated that ambient noise structure differs fundamentally between plate-based and droplet-based platforms. Cargnelli et al. (12) noted scAR’s negative impact on batch integration and distortion of negative control datasets, but did not demonstrate cell type emergence directly. Notably, Cargnelli et al. (12) were unable to execute scAR in full mode due to computational resource constraints and evaluated it in reduced mode only, which may affect the comparability of scAR results between the two benchmarks. The present data establish that scAR and CellClear do not merely over-correct but introduce biologically unsubstantiated signal with direct implications for any study in which ambient correction precedes cell type annotation, trajectory inference, or rare population identification.

### Reliable performance: CellBender and SoupX

Among the four tools that did not exhibit count inflation, CellBender consistently achieved the highest sensitivity and precision across the five hgmm datasets included in aggregate analyses (Figure 2E). SoupX followed as the most computationally lightweight option, offering robust removal with minimal count distortion and the fastest runtime of any tool evaluated on the largest dataset (Figure 5A, 5C). This performance ranking is consistent across three independent benchmarks employing different ground truth strategies: genotype-mixing in mouse kidney (3), species-mixing in hgmm datasets (this study), and simulation combined with genotype-mixing (12). Convergence across methodologically distinct approaches strengthens confidence in this ordering.

### Impact on complex tissue annotation

Beyond contamination removal metrics, ambient correction produced measurable but heterogeneous improvements in complex tissue datasets.:

Marker gene enrichment scores improved significantly following correction with CellBender, SoupX, and DecontX across all three complex tissue datasets (Figure 3A). DecontX achieved the largest median enrichment changes despite its lower removal sensitivity in the hgmm benchmarks. This apparent paradox is mechanistically consistent: the marker gene enrichment metric measures the relative specificity of canonical markers in their expected cell type versus all other cells, and a tool that removes contamination conservatively while preserving endogenous gene expression can improve marker specificity at least as effectively as a more aggressive tool, provided the contamination it does remove includes the cross-cell-type ambient signal that suppresses marker ratios. The result suggests that for annotation-focused applications, conservative high-specificity removal may be preferable to aggressive removal, and that sensitivity alone is an incomplete performance criterion even in the absence of count inflation.

The recovery of Platelets in WBC data represents the clearest demonstration of ambient correction utility in this benchmark. This population was absent from confidently annotated cells in uncorrected data but was recovered by every globally performing ambient removal tool evaluated - including tools with otherwise very different correction scopes and algorithms. The convergence across tools with distinct architectures (global VAE, Bayesian mixture, maximum likelihood subtraction) argues against any single tool-specific artifact and instead establishes that Platelet marker suppression in uncorrected data reflects genuine ambient RNA interference with canonical platelet transcripts (PF4, PPBP) rather than true biological absence. This is precisely the use case ambient correction tools were designed for: a low-abundance population whose defining transcripts are co-expressed by or derived from abundant contaminating cell types. However, both tools utilizing gene-selective ambient removal, namely CellClear and scCDC, were either not applicable to well-plate based datasets (CellClear) or did not show the recovery of platelets (scCDC) despite otherwise successful application.

Gains in annotation confidence were substantially larger in the WBC and PFC datasets than in PBMC, where baseline annotation quality was near-perfect. This pattern suggests the practical benefit of ambient correction scales with contamination severity and baseline annotation difficulty, and that already high-quality datasets may gain little in annotation terms while remaining exposed to the count distortion risks demonstrated for scAR and CellClear.

### Platform scope: DecontX and the well-plate finding

DecontX occupies a distinct operational niche among the tools evaluated. It is the only tool in this panel that both operates without a raw unfiltered count matrix and is not restricted to droplet-based platforms, properties that follow from its cluster-based Bayesian mixture model estimating the ambient profile from cell-containing droplets rather than empty ones (4). scCDC shares the same properties but was not validated on well-plate based datasets and further shows worse sensitivity and precision when compared to DecontX. This makes DecontX the only practical option in the common scenario where only a filtered count matrix is available, as is frequently the case for publicly deposited datasets, and the only tool whose input assumptions are not violated on well-plate-based platforms. In the BD Rhapsody WBC dataset, DecontX is accordingly the only tool in this panel whose input assumptions are not violated regardless of data availability (Figure 5B), a finding with practical relevance given the growing use of well-plate-based and microfluidic platforms, including BD Rhapsody, MARS-seq, and Smart-seq variants, particularly in immunological and clinical settings where throughput, cost, or sample type constraints favor non-droplet approaches (6). DecontX’s performance on the hgmm metrics ranks mid-range with lower removal sensitivity than CellBender (Figure 2C, 2E), consistent with Cargnelli et al.’s (12) observation that DecontX preserves endogenous RNA well at the cost of incomplete ambient removal. Importantly, the present data suggest that access to the raw count matrix is not a prerequisite for meaningful DecontX correction in the well-plate context: enrichment outcomes for DecontX° (without background) and DecontX (with background) did not differ significantly in the WBC dataset (pairwise Wilcoxon, p = 0.074), and confidence score improvements were comparable across annotation models, although performance of DecontX without raw matrix input on droplet-based hgmm datasets was inferior (data not shown), suggesting this finding may be platform-context-dependent. This finding has direct practical relevance for the common scenario of re-analyzing publicly deposited data without accompanying raw count matrices, and argues that DecontX should be considered even when only a filtered count matrix is available. The ambient noise structure in well-plate platforms differs physically from droplet-based systems, arising in part through cross-well contamination rather than droplet co-encapsulation (Colino-Sanguino et al., 2024), and whether any tool validated primarily on droplet data is fully appropriate for plate-based platforms remains an open question that a dedicated benchmark with appropriate ground truth should address.

### Scalability and computational constraints

The order-of-magnitude runtime differences documented in Figure 5A have practical implications beyond computational convenience. CellClear’s failure to complete within feasible time on the 172,000-nucleus PFC dataset disqualifies it for atlas-scale applications irrespective of its correction quality, unraveling a constraint that is not apparent from benchmarks conducted exclusively on smaller datasets. CellBender’s scaling behavior on large inputs, while partially an artifact of the modest GPU used here, reflects genuine architectural costs of the variational autoencoder approach that users should account for when planning large cohort analyses. In contrast, the SoupX alternative incurs none of these costs. DecontX’s disproportionate runtime on the PFC dataset, attributed to memory pressure rather than algorithmic scaling, suggests that its practical upper limit may be closer to 100,000 cells than the current benchmark implies, and warrants profiling on datasets of intermediate size before deployment in large studies. However, relative comparisons between tools remain informative, but absolute values should be interpreted accordingly.

### Limitations

Several limitations of this benchmark should be stated directly. The hgmm ground truth is partial: because intra-species ambient counts (human-in-human, mouse-in-mouse) cannot be directly observed, expected intra-species removal was extrapolated by assuming that human ambient RNA contaminates human cells at the same rate as it contaminates mouse cells, and equivalently for mouse ambient RNA in human cells. This assumes equal encapsulation probability across cell types, which is biologically reasonable given the stochastic nature of droplet capture but cannot be directly verified. Consequently, precision and specificity values should be interpreted as approximations rather than exact estimates. The 1k dataset was excluded from aggregate metrics due to its markedly lower ambient contamination, and contamination-level-dependent performance variation across the full range deserves dedicated study. Only one well-plate dataset was analyzed, and generalizability across BD Rhapsody library types, other well-plate platforms, and non-immune tissue types remains to be established. Some tools, particularly SoupX, offer user-tunable parameters that could meaningfully shift results for specific datasets; benchmarking defaults does not capture this flexibility. GPU benchmarking was performed on consumer-grade hardware, and CellBender and scAR runtimes at large scale should be considered upper bounds rather than representative values. Finally, FastCAR (11) was excluded on methodological grounds, as it applies gene-level threshold subtraction optimized for DGE analysis rather than global ambient removal, making it incompatible with the global removal metrics employed here. scCDC (9) was included as a reference comparator with the explicit caveat that its gene-selective correction paradigm means sensitivity and precision metrics reflect a narrower correction target than the other tools; direct comparison of its removal metrics with those of global correction tools should be interpreted accordingly. A future benchmark should accommodate gene-selective and DGE-optimized correction approaches as fully distinct evaluation categories with appropriate metrics. Further, the marker gene enrichment analyses are based on n=9 markers per dataset, and the pairwise Wilcoxon signed-rank tests comparing hgmm sensitivity across tools are limited to n=5 paired observations, for which the minimum achievable p-value is 0.0313. Results at this boundary should be interpreted as the maximum achievable significance level at this sample size rather than strong evidence of effect magnitude. Conclusions drawn from these tests are accordingly treated as directional and consistent with, rather than independently establishing, the quantitative performance ordering.

### Practical recommendations and outlook

As ambient RNA removal becomes embedded in standardized pipelines and applied as a default step without prior evidence of contamination (12), the risk of introducing model-driven artifacts grows, as demonstrated in the in-depth analyses of corrected scAR and CellClear matrices. At the same time, adoption of ambient correction tools remains low relative to overall sxRNA-seq usage (Figure 1B), and the present benchmark argues for broader uptake: for the three tools recommended here, the risk of harm is low (no artificial count introduction, high specificity) and the potential to improve marker gene specificity, annotation confidence, and recovery of low-abundance populations is real, particularly in lower-quality / highly contaminated datasets. The primary message is not caution but informed selection. The tools exist, they work, and with appropriate choice they improve data quality.

## Conclusion

This benchmark establishes count matrix integrity, i.e. the absence of corrected values exceeding their raw counterparts, as an indispensable criterion for ambient RNA correction tool evaluation, alongside removal sensitivity. scAR and CellClear both fail this criterion: scAR introduces artificial counts at 17-60% of total raw counts across datasets and generated up to eight spurious cell types in automated annotation, while CellClear replaces more than 93% of the original count matrix with values derived from matrix factorization. The fact that tools achieving competitive sensitivity scores on standard ground-truth benchmarks can simultaneously restructure the count matrix in biologically unsubstantiated ways reveals a systematic blind spot in how ambient correction tools are currently evaluated and reported. We recommend that count inflation quantification be adopted as a routine validation step in both tool development and benchmarking.

Among tools that do not exhibit count inflation, CellBender provides the highest sensitivity and precision for droplet-based data; SoupX offers a robust and computationally lightweight alternative. On non-droplet platforms and for datasets lacking raw count matrices - both common scenarios in the reanalysis of publicly deposited data - DecontX is the only tool in this panel whose input assumptions are not violated.It produced substantive annotation improvements in the BD Rhapsody WBC dataset though generalizability across plate-based platforms and tissue types warrants dedicated evaluation. The convergence of CellBender, SoupX, and DecontX as the recommended tools across three independent benchmarks employing distinct ground truth strategies provides an unusually strong basis for practical guidance. Tool selection should be governed by platform compatibility and count integrity first, with sensitivity as a secondary discriminator among integrity-passing tools.

## Methods

### Datasets

Six publicly available human-mouse cell line mixture (hgmm) datasets from 10x Genomics were used to benchmark ambient RNA removal performance against a partial ground truth (Table 2). All datasets consist of HEK293T (human) and NIH3T3 (mouse) cells sequenced on the 10x Chromium platform and were obtained as CellRanger-processed filtered and raw feature-barcode matrices directly from the 10x Genomics website. Dataset names reflect the nominal cell count targets; actual post-QC cell numbers are reported in Table 2.

**Table 2:**
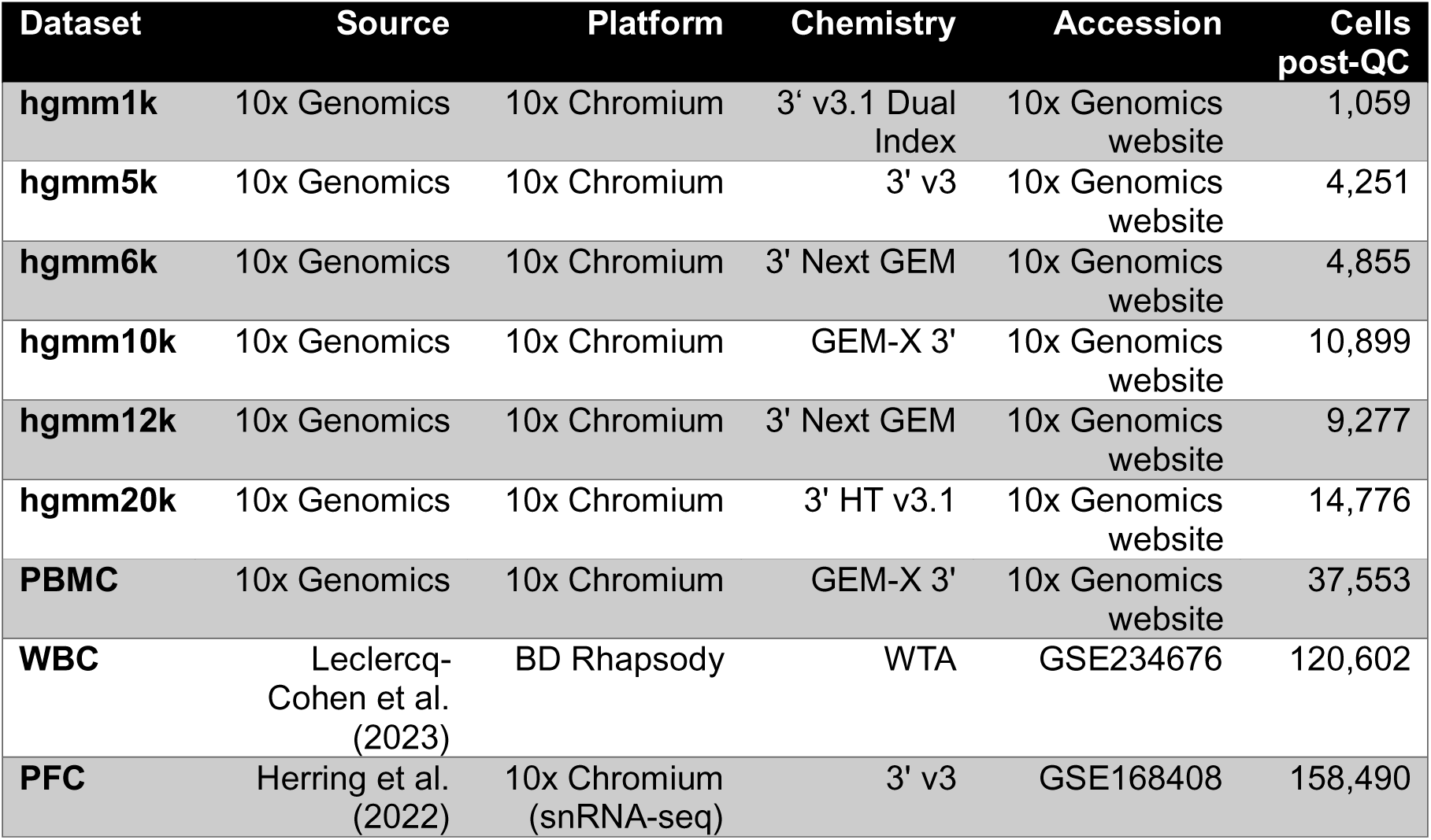
Overview of datasets used in this benchmark. Datasets used in this benchmark. Six human-mouse cell line mixture (hgmm) datasets from 10x Genomics provide partial ground truth for quantitative performance evaluation. Three complex tissue datasets from published studies and public repositories were used to assess tool behavior under biologically relevant conditions. Cells post-QC reflects the number of cells retained after quality control filtering; for CellBender and CellClear, which operate on unfiltered raw input and apply QC post-correction, final cell counts per tool may differ marginally from values reported here.

| Dataset | Source | Platform | Chemistry | Accession | Cells post-QC |
| --- | --- | --- | --- | --- | --- |
| hgmm1k | 10x Genomics | 10x Chromium | 3' v3.1 Dual Index | 10x Genomics website | 1,059 |
| hgmm5k | 10x Genomics | 10x Chromium | 3' v3 | 10x Genomics website | 4,251 |
| hgmm6k | 10x Genomics | 10x Chromium | 3' Next GEM | 10x Genomics website | 4,855 |
| hgmm10k | 10x Genomics | 10x Chromium | GEM-X 3' | 10x Genomics website | 10,899 |
| hgmm12k | 10x Genomics | 10x Chromium | 3' Next GEM | 10x Genomics website | 9,277 |
| hgmm20k | 10x Genomics | 10x Chromium | 3' HT v3.1 | 10x Genomics website | 14,776 |
| PBMC | 10x Genomics | 10x Chromium | GEM-X 3' | 10x Genomics website | 37,553 |
| WBC | Leclercq-Cohen et al. (2023) | BD Rhapsody | WTA | GSE234676 | 120,602 |
| PFC | Herring et al. (2022) | 10x Chromium (snRNA-seq) | 3' v3 | GSE168408 | 158,490 |

Three complex tissue datasets were used to evaluate tool performance under biologically relevant conditions. A peripheral blood mononuclear cell (PBMC) dataset (10x Genomics: “60k Human PBMCs Stained with 8 TotalSeq-B Human Hashtags”, 4 donors; 37,553 cells post-QC) was selected as a droplet-based immune tissue reference. A human whole blood leukocyte (WBC) dataset (PMID: 37379429; 120,602 cells post-QC) (15) generated on the BD Rhapsody well-plate platform was included as the sole non-droplet dataset, representing a platform category not evaluated by prior benchmarks. A human prefrontal cortex (PFC) single-nucleus RNA-seq dataset (16) (PMID: 36318921; 158,490 nuclei post-QC) was included as a neuronal tissue reference.

### Data Processing

All datasets were processed using a standardized pipeline implemented in Python 3.12.9 with Scanpy 1.11.1 (17). Quality control was performed using automated thresholding based on the Median Absolute Deviation (MAD), adapted from Heumos et al. (18). Cells deviating more than 5 MADs in total count number, number of detected genes, or percentage of counts in the 20 most highly expressed genes were excluded. Additionally, cells with mitochondrial count fractions exceeding 8% (PBMC, PFC, hgmm1k; 10k) or 20% (hgmm5k;6k;12k;20k, WBC) were removed, applying stringent and permissive thresholds to high- and lower-quality datasets respectively. For the WBC and PFC datasets, which comprise multiple batches, MAD thresholds were computed per sample.

Doublets were detected using the Scrublet (19) implementation in Scanpy 1.11.1 (17) and excluded prior to downstream analysis. For multi-sample datasets (WBC, PFC), doublet scoring was performed in a batch-aware manner. No batch effect correction was applied: in the PFC dataset, each sample constitutes a biological replicate and correction risked masking true biological variation; in the WBC dataset, batch correction was deliberately omitted to preserve the detectability of ambient RNA correction effects on dimensionality reduction and clustering.

Ambient RNA correction was applied at the position in the pipeline appropriate to each tool’s input requirements (see Tool application). Following correction, counts were normalized to the per-cell median library size and log1p-transformed. PCA was computed on z-standardized counts (clipped to ±10 SD); UMAP and a k-nearest-neighbor graph (default settings) were computed on the principal components. Clustering was performed with the Leiden algorithm (resolution 0.5 and 1.0). Feature selection (highly variable genes) was intentionally omitted to avoid masking tool-specific effects on the count matrix structure.

### Species classification (hgmm)

Human and mouse cells were distinguished using a count-ratio classifier. For each cell, the cross-species count ratio was defined as the fraction of total counts originating from the non-majority species. Cells were assigned a preliminary species label based on which species contributed the majority of counts. A dataset-specific classification threshold was then derived from the 5th percentile of the cross-species count ratio distribution within each majority-label group, except for the 1k dataset where the 1st percentile was used due to a markedly narrower distribution of non-majority counts. Cells with cross-species ratios exceeding this threshold were classified as ambiguous and excluded from all downstream analyses, as they likely represent inter-species technical doublets. This dynamic thresholding approach yielded an effective ambiguity boundary near 12% cross-species counts across most datasets.

### Tool application

All six tools were applied using their respective default parameters unless otherwise noted. Tool versions were: CellBender 0.3.0, DecontX (celda v1.4.1), SoupX 1.6.2, scCDC 1.4, scAR 0.7.0, CellClear 0.0.3.

CellBender was applied directly to the raw (unfiltered) count matrix via the remove-background command-line function. The parameters expected-cells and total-droplets-included were estimated individually per dataset from barcode rank plots; for the PBMC dataset, CellBender default parameter estimation was used. For all other datasets, values were set as follows: hgmm1k (2,000 / 10,000), hgmm5k (5,000 / 30,000), hgmm6k (6,000 / 25,000), hgmm10k (13,000 / 40,000), hgmm12k (12,000 / 30,000), hgmm20k (15,000 / 50,000), WBC (120,000 / 132,000), PFC (170,000 / 300,000). All CellBender runs used 150 training epochs.

DecontX was run directly in R on filtered, unnormalized count matrices with the raw (unfiltered) matrix provided as background for empirical contamination profile estimation (background mode). Cluster labels were inferred internally using the default UMAP/dbscan approach. DecontX was additionally run without providing the raw unfiltered count matrix (denoted DecontX°) for the well-plate-based WBC dataset to assess correction capacity under the common scenario where only filtered count matrices are publicly available.

SoupX was run in Python using rpy2 with cluster labels derived from Leiden clustering (resolution 1.0) computed in Scanpy on pre-QC filtered counts. The contamination fraction was estimated automatically using autoEstCont and counts were corrected using adjustCounts with default parameters.

scCDC was applied via its default pipeline to QC-filtered count matrices without providing empty droplet data, consistent with its design requirements. Prior to correction, count matrices were processed within R following scCDC’s recommended workflow, including normalization, scaling, PCA, and Louvain clustering (resolution = 0.5). This preprocessing is required for scCDC’s contamination-causing gene detection step. The corrected count matrices output by scCDC represent adjusted raw-scale counts and were subsequently reimported into the Python pipeline for downstream analysis.

scAR was run using the setup_anndata interface with recommended settings. The ambient RNA profile was estimated from cell-free droplets and the model trained for 200 epochs on filtered counts. For the WBC (BD Rhapsody) dataset, the prob parameter was set to 0.95 instead of the default 0.995 to accommodate the well-plate platform format.

CellClear was applied via the correct_expression command-line function using both raw and filtered files as input and default parameters. Note that CellBender and CellClear require quality control to be applied post-correction, as they operate on raw or pre-QC input; all other tools received QC-filtered matrices as input.

### Benchmarking metrics

For each hgmm dataset and tool, per-cell count totals were aggregated into four categories: human counts in human cells (hg-hg), mouse counts in mouse cells (mm-mm), mouse counts in human cells (mm-hg), and human counts in mouse cells (hg-mm). The latter two constitute the directly observable inter-species ambient counts.

**Sensitivity** was defined as the percentage of total known inter-species ambient counts removed after correction:

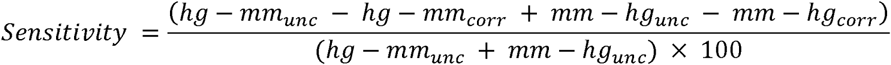

**Precision (extrapolated):** Because total count removal includes both inter- and intra-species counts, a naive precision (inter-species removed / total removed) would underestimate true precision by ignoring legitimate intra-species ambient removal. Expected intra-species ambient counts were extrapolated from the observed inter-species ratios: expected_hg_ambient = (hg-in-mm / total_mouse) × total_human, and analogously for mouse. Intra-species true positives were then estimated per species separately, capping each at its respective expected ambient: intra_TP_hg = min(hg_removed, expected_hg_ambient), and analogously for mouse. This per-species capping ensures that over-removal in one species cannot be masked by proportionate under-removal in the other. Extrapolated precision was then:

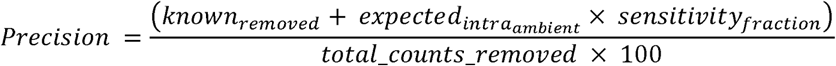

capped at 100%.

**Specificity** was computed from a per-species confusion matrix framework. False positives were defined separately for each species as intra-species count removal exceeding the expected intra-species ambient (FP_hg = max(0, hg_removed − expected_hg_ambient); analogously for mouse) and summed across species. Inter-species removal is excluded from FP entirely, as any hg-in-mm or mm-in-hg removal is by definition a true positive. True negatives were computed as TN = (total_counts_unc − expected_ambient) − FP, where expected_ambient is the sum of known inter-species and extrapolated intra-species ambient. Specificity was then:

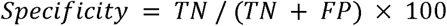

**Intra-species removal ratio** was defined as the geometric mean of the human and mouse intra-species removal ratios: observed intra-species counts removed divided by the expected intra-species ambient counts derived above. A ratio of 1.0 is the expected value under the assumption that a tool removes intra-species ambient proportionally to its observed inter-species sensitivity, i.e. it does not indicate perfect specificity, but rather internally consistent behavior. Values substantially above 1.0 indicate removal beyond what ambient RNA alone can account for, i.e. non-specific removal of endogenous counts.

**Consistency** was defined as 100 minus the coefficient of variation (CV) of per-dataset sensitivity, computed across the five hgmm datasets included in aggregate analyses. The 1k dataset was excluded from all aggregate metrics due to markedly lower baseline inter-species contamination relative to all larger datasets, which would inflate apparent variability independent of tool behavior.

### Count inflation analysis

To quantify artificial count inflation, we decomposed corrected count matrices into removed counts (corrected < raw) and added counts (corrected > raw) on a per-cell, per-gene basis. The fraction of total counts removed and artificially added relative to the original raw total were calculated for each tool and dataset. To detect spurious cell type emergence, we compared CellTypist annotations between uncorrected and corrected data; cell types present post-correction but absent in uncorrected data were classified as “emerged.” For each emerged cell type, we computed a zero-to-positive (0→+) inflation metric: the proportion of cells in which a given gene had zero raw counts but non-zero corrected counts and reported the gene with the highest such proportion alongside its mean corrected expression.

### Complex tissue analysis

**Cell type annotation:** Following ambient RNA correction and the data processing steps described above, automated cell type annotation was performed using CellTypist 1.6.3 with majority voting enabled. The Immune_All_Low and Immune_All_High models were applied to both the PBMC and WBC datasets; the Adult_Human_PrefrontalCortex model was used for the PFC dataset. No cell type annotation was performed on hgmm datasets.

**Marker gene enrichment analysis:** For each complex tissue dataset, marker genes were selected from CellMarker 2.0 and validated by inspecting average expression across annotated cell types (Supplementary Figure 4). The log□-fold enrichment ratio change was calculated as log□[(x_corr/y_corr) / (x_unc/y_unc)], where x denotes marker gene counts in the expected cell type and y denotes counts in all other cells, for corrected and uncorrected data respectively. scAR and CellClear were excluded from statistical testing due to systematic count inflation invalidating ratio-based comparisons. Among the remaining tools, differences in enrichment distributions were assessed using a Friedman test across markers (treating each marker as a repeated measure). Post-hoc one-sided Wilcoxon signed-rank tests against zero (H□: median enrichment change > 0) were applied when the Friedman test indicated significance (α = 0.05), with Benjamini-Hochberg correction across tools. Pairwise Wilcoxon signed-rank tests between tools were additionally computed.

**Confidence score analysis:** CellTypist confidence scores were compared between corrected and uncorrected conditions using two-sided Mann-Whitney U tests, with Bonferroni correction across tools within each dataset. Median differences relative to uncorrected baseline were additionally reported as a measure of practical significance.

**Cell type abundance changes:** The number of confidently annotated cells per cell type was recorded for each tool and compared to the uncorrected baseline, using CellTypist confidence score thresholds of > 0.9 for coarse annotation models and > 0.5 for fine annotation models, reflecting the higher per-class specificity of coarse models. Cell types absent in the uncorrected data but detected after correction were classified as “Gain”; cell types present in uncorrected data but absent after correction were classified as “Loss”. Remaining differences were expressed as percentage change relative to uncorrected counts.

### Runtime benchmarking

Runtimes were measured as wall-clock time encompassing input file reading, ambient RNA correction, and output writing. All benchmarks were run on an Intel Core i5 processor with 32 GB RAM and 6 cores. For machine learning-based tools (CellBender, scAR), runtimes were additionally measured on an NVIDIA GeForce GTX 1050 Ti (4 GB VRAM, 768 CUDA cores). Runtimes were evaluated across datasets of increasing cell number (hgmm20k with ∼18,000 cells - PFC dataset with ∼ 172,000 cells post-QC) to characterize scaling behavior.

### Summary scoring / radar plots

Radar plot scores were computed as follows. Sensitivity, Specificity, Precision, and Consistency were derived from quantitative metrics described in Figure 2/3 methods. Speed was computed using log-scale min-max normalization: 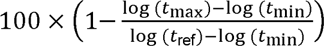 where t_max_ is the tool’s maximum observed runtime, t_min_ is the minimum across all tools (SoupX, 6.5 h), and t_ref_ is the maximum across all tools (CellClear, 582 h). GPU runtimes were used where available. Log scaling was applied to account for the order-of-magnitude differences in runtime and to prevent CellClear’s outlier performance from artificially compressing scores of all other tools into a narrow band. Versatility was assigned based on operational criteria: DecontX and scCDC received the maximum score (100) as they run without raw count matrices and are applicable to non-droplet platforms. CellBender and SoupX received 60 reflecting droplet-platform dependency and raw data requirements; scAR and CellClear received 20 and 10, respectively, additionally penalized for artificial count inflation that critically limits the utility of their output.

### Citation analysis

We assessed the adoption of ambient RNA removal tools relative to overall scRNA-seq usage through PubMed citation analysis. Using the NCBI Entrez E-utilities API (Biopython v1.86), we retrieved all papers citing Seurat (versions 1-5, PMIDs: 25867923, 29608179, 31178118, 34062119, 37231261) and Scanpy (PMID: 29409532) as a proxy for scRNA-seq analysis papers. To enrich for papers reporting empirical analyses, we further required citing papers to have an associated GEO dataset entry, as data deposition is standard practice for studies generating or reanalysing sequencing data. This GEO-filtered set (n=8,682) served as the denominator. Within this set, we identified papers additionally citing SoupX (PMID: 33367645), DecontX (PMID: 32138770), CellBender (PMID: 37550580), FastCAR (PMID: 38030970), or scCDC (PMID: 38783325). For tools with multiple versions (Seurat), citing papers were deduplicated such that a paper citing multiple versions was counted once. Papers citing multiple tools within the same category were counted once per tool; overlap was modest (6.8% for Seurat/Scanpy, 3.3% for ambient tools). Citation counts were stratified by publication year of the citing paper. Remaining limitations include studies depositing data to non-GEO repositories (e.g. ArrayExpress) being excluded from the denominator, and potential incomplete coverage of PubMed citation linkages. This citation analysis was conducted in February 2026.

### Software and code availability

Ambient RNA correction tools were applied at the following versions: CellBender 0.3.0, SoupX 1.6.2, DecontX 1.4.1 (celda), scCDC 1.4, scAR 0.7.0, and CellClear 0.0.3. All Python-based analyses were performed with Python 3.12.9 using the following packages: Scanpy 1.11.1, NumPy 2.1.3, pandas 2.2.3, matplotlib 3.10.1, seaborn 0.13.2, anndata 0.11.4, CellTypist 1.6.3, and rpy2 3.5.17. DecontX was run in R 4.4.3; conversion between AnnData and SingleCellExperiment objects was performed using zellkonverter 1.16.0. All analysis code is available under github at https://github.com/luschroed/Benchmarking and further released at https://doi.org/10.5281/zenodo.19348468.

Large language model (LLM) tools were used for manuscript language editing and for assistance with visualization code. No LLM was used for data analysis, statistical inference, or interpretation of results.

## Declarations

### Ethics approval and consent to participate

Not applicable.

### Consent for publication

Not applicable.

### Availability of data and materials

All datasets analyzed during this study are publicly available. hgmm datasets were obtained from the 10x Genomics website. PBMC data were obtained from the 10x Genomics website. WBC data are available under GEO accession GSE234676. PFC data are available under GEO accession GSE168408. All analysis code is available at https://github.com/luschroed/Benchmarking [https://doi.org/10.5281/zenodo.19367740].

### Authors’ contributions

**LS:** Investigation, Software, Data curation, Validation, Writing - original draft, Writing - review & editing

**NR:** Conceptualization, Methodology, Formal analysis, Visualization, Writing - original draft, Writing - review & editing, Project administration, Supervision

**SG:** Supervision, Writing - review & editing

## Bibliography

1. Young MD, Behjati S. SoupX removes ambient RNA contamination from droplet-based single-cell RNA sequencing data. GigaScience. 2020 Dec 26;9(12):giaa151. doi:10.1093/gigascience/giaa151 PubMed PMID: 33367645; PubMed Central PMCID: PMC7763177.

2. Fleming SJ, Chaffin MD, Arduini A, Akkad AD, Banks E, Marioni JC, et al. Unsupervised removal of systematic background noise from droplet-based single-cell experiments using CellBender. Nat Methods. 2023 Sep;20(9):1323–35. doi:10.1038/s41592-023-01943-7 PubMed PMID: 37550580.

3. Janssen P, Kliesmete Z, Vieth B, Adiconis X, Simmons S, Marshall J, et al. The effect of background noise and its removal on the analysis of single-cell expression data. Genome Biol. 2023 Jun 19;24(1):140. doi:10.1186/s13059-023-02978-x PubMed PMID: 37337297; PubMed Central PMCID: PMC10278251.

4. Yang S, Corbett SE, Koga Y, Wang Z, Johnson WE, Yajima M, et al. Decontamination of ambient RNA in single-cell RNA-seq with DecontX. Genome Biol. 2020 Mar 5;21(1):57. doi:10.1186/s13059-020-1950-6 PubMed PMID: 32138770; PubMed Central PMCID: PMC7059395.

5. Arceneaux D, Chen Z, Simmons AJ, Heiser CN, Southard-Smith AN, Brenan MJ, et al. A contamination focused approach for optimizing the single-cell RNA-seq experiment. iScience. 2023 Jun 29;26(7):107242. doi:10.1016/j.isci.2023.107242 PubMed PMID: 37496679; PubMed Central PMCID: PMC10366499.

6. Colino-Sanguino Y, Fuente LR de la, Gloss B, Law AMK, Handler K, Pajic M, et al. Performance comparison of high throughput single-cell RNA-Seq platforms in complex tissues. Heliyon. 2024 Sep 15;10(17). doi:10.1016/j.heliyon.2024.e37185 PubMed PMID: 39296129.

7. Ewels PA, Peltzer A, Fillinger S, Patel H, Alneberg J, Wilm A, et al. The nf-core framework for community-curated bioinformatics pipelines. Nat Biotechnol. 2020 Mar;38(3):276–8. doi:10.1038/s41587-020-0439-x

8. Sheng C, Lopes R, Li G, Schuierer S, Waldt A, Cuttat R, et al. Probabilistic machine learning ensures accurate ambient denoising in droplet-based single-cell omics [Internet]. bioRxiv; 2022 [cited 2026 Mar 3]. p. 2022.01.14.476312. Available from: https://www.biorxiv.org/content/10.1101/2022.01.14.476312v4 doi:10.1101/2022.01.14.476312

9. Wang W, Cen Y, Lu Z, Xu Y, Sun T, Xiao Y, et al. scCDC: a computational method for gene-specific contamination detection and correction in single-cell and single-nucleus RNA-seq data. Genome Biol. 2024 May 23;25:136. doi:10.1186/s13059-024-03284-w PubMed PMID: 38783325; PubMed Central PMCID: PMC11112958.

10. Huang W, Hu L, Qian Z, Fan J. CellClear: Enhancing Single-cell RNA Data Quality via Biologically-Informed Ambient RNA Correction [Internet]. bioRxiv; 2025 [cited 2026 Mar 3]. p. 2024.08.05.606571. Available from: https://www.biorxiv.org/content/10.1101/2024.08.05.606571v2 doi:10.1101/2024.08.05.606571

11. Berg M, Petoukhov I, van den Ende I, Meyer KB, Guryev V, Vonk JM, et al. FastCAR: fast correction for ambient RNA to facilitate differential gene expression analysis in single-cell RNA-sequencing datasets. BMC Genomics. 2023 Nov 29;24(1):722. doi:10.1186/s12864-023-09822-3 PubMed PMID: 38030970; PubMed Central PMCID: PMC10687889.

12. Cargnelli CB, Nielsen JV, Madsen JGS. Benchmarking computational decontamination of ambient RNA [Internet]. bioRxiv; 2026 [cited 2026 Mar 3]. p. 2026.01.13.699237. Available from: https://www.biorxiv.org/content/10.64898/2026.01.13.699237v1 doi:10.64898/2026.01.13.699237

13. Domínguez Conde C, Xu C, Jarvis LB, Rainbow DB, Wells SB, Gomes T, et al. Cross-tissue immune cell analysis reveals tissue-specific features in humans. Science. 2022 May 13;376(6594):eabl5197. doi:10.1126/science.abl5197

14. Caglayan E, Liu Y, Konopka G. Neuronal ambient RNA contamination causes misinterpreted and masked cell types in brain single-nuclei datasets. Neuron. 2022 Dec 21;110(24):4043–4056.e5. doi:10.1016/j.neuron.2022.09.010 PubMed PMID: 36240767; PubMed Central PMCID: PMC9789184.

15. Leclercq-Cohen G, Steinhoff N, Albertí Servera L, Nassiri S, Danilin S, Piccione E, et al. Dissecting the Mechanisms Underlying the Cytokine Release Syndrome (CRS) Mediated by T-Cell Bispecific Antibodies. Clin Cancer Res Off J Am Assoc Cancer Res. 2023 Nov 1;29(21):4449–63. doi:10.1158/1078-0432.CCR-22-3667 PubMed PMID: 37379429; PubMed Central PMCID: PMC10618647.

16. Herring CA, Simmons RK, Freytag S, Poppe D, Moffet JJD, Pflueger J, et al. Human prefrontal cortex gene regulatory dynamics from gestation to adulthood at single-cell resolution. Cell. 2022 Nov 10;185(23):4428–4447.e28. doi:10.1016/j.cell.2022.09.039

17. Wolf FA, Angerer P, Theis FJ. SCANPY: large-scale single-cell gene expression data analysis. Genome Biol. 2018 Feb 6;19(1):15. doi:10.1186/s13059-017-1382-0 PubMed PMID: 29409532; PubMed Central PMCID: PMC5802054.

18. Heumos L, Schaar AC, Lance C, Litinetskaya A, Drost F, Zappia L, et al. Best practices for single-cell analysis across modalities. Nat Rev Genet. 2023 Aug;24(8):550–72. doi:10.1038/s41576-023-00586-w

19. Wolock SL, Lopez R, Klein AM. Scrublet: Computational Identification of Cell Doublets in Single-Cell Transcriptomic Data. Cell Syst. 2019 Apr 24;8(4):281–291.e9. doi:10.1016/j.cels.2018.11.005

