## Supplementary Data for "Benchmarking ambient RNA removal across droplet and well-plate platforms reveals artificial count generation as a critical failure mode of scAR and CellClear"

### Supplementary Figures

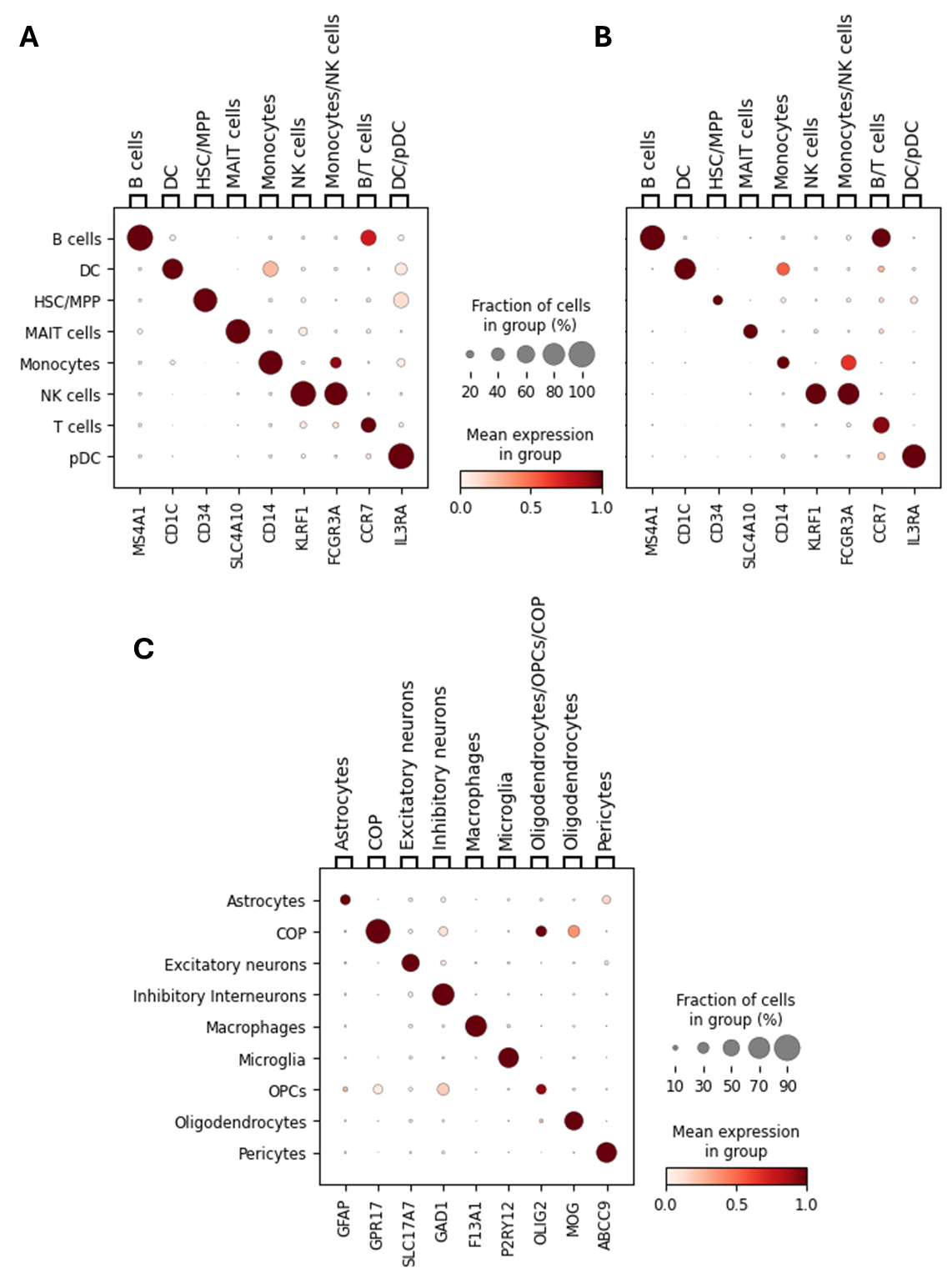

Supplementary Figure 1: Selected marker genes based on CellMarker 2.0

Dot plots representing the mean expression of each marker gene in the detected cell types without ambient RNA correction in the **(A)** PBMC, **(B)** WBC, and **(C)** PFC dataset.

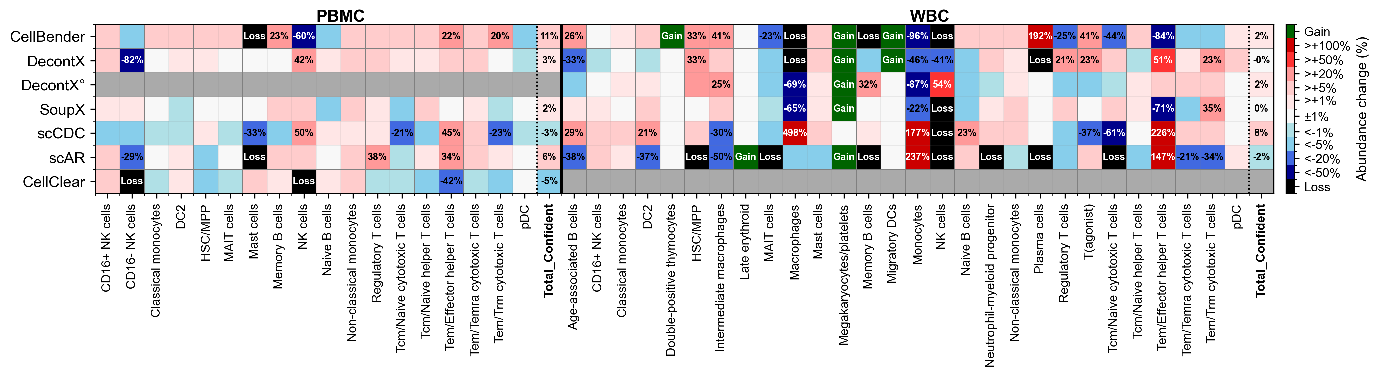

Supplementary Figure 2: Cell type abundance heatmap, fine annotation model, PBMC and WBC

Heatmap of relative cell type abundance changes after ambient RNA correction compared to uncorrected counts, as in Figure 3C, but using the fine-resolution CellTypist annotation models (Immune_All_Low for PBMC; Immune_All_High for WBC). Gains (newly detected cell types) and losses (complete disappearance) are annotated explicitly; percentage changes >20% or for the Total_Confident summary column are labeled. Grey cells indicate tool–dataset combinations not applicable. scAR and CellClear shown for reference but excluded from statistical inference due to count inflation artifacts. Bold column labels denote the Total_Confident summary column.

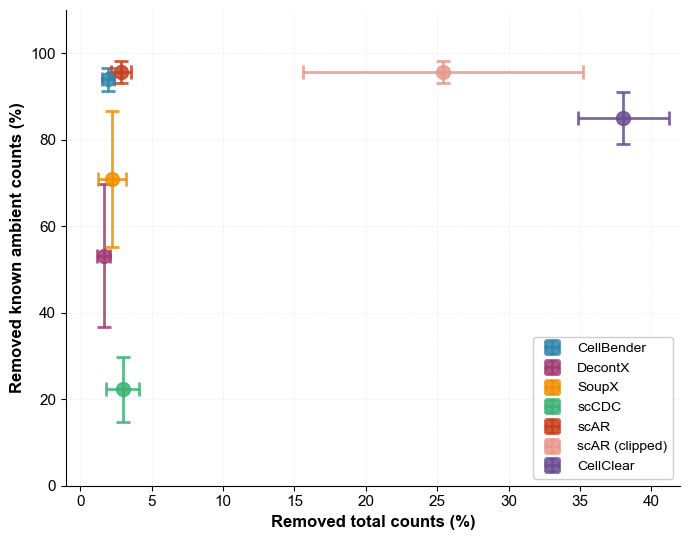

Supplementary Figure 3: Figure 2B variant with scAR counts clipped to raw maximum

Intra-species removal ratios (observed/expected) as in Figure 2B, with scAR corrected counts clipped per cell and gene to not exceed the corresponding raw count value prior to metric calculation. Clipping eliminates the artificial count inflation documented in Figure 4 and shows where scAR would place on the intra-species metric under a count-conservative assumption. CellClear excluded (ratio 15–45×, as in Figure 2B).

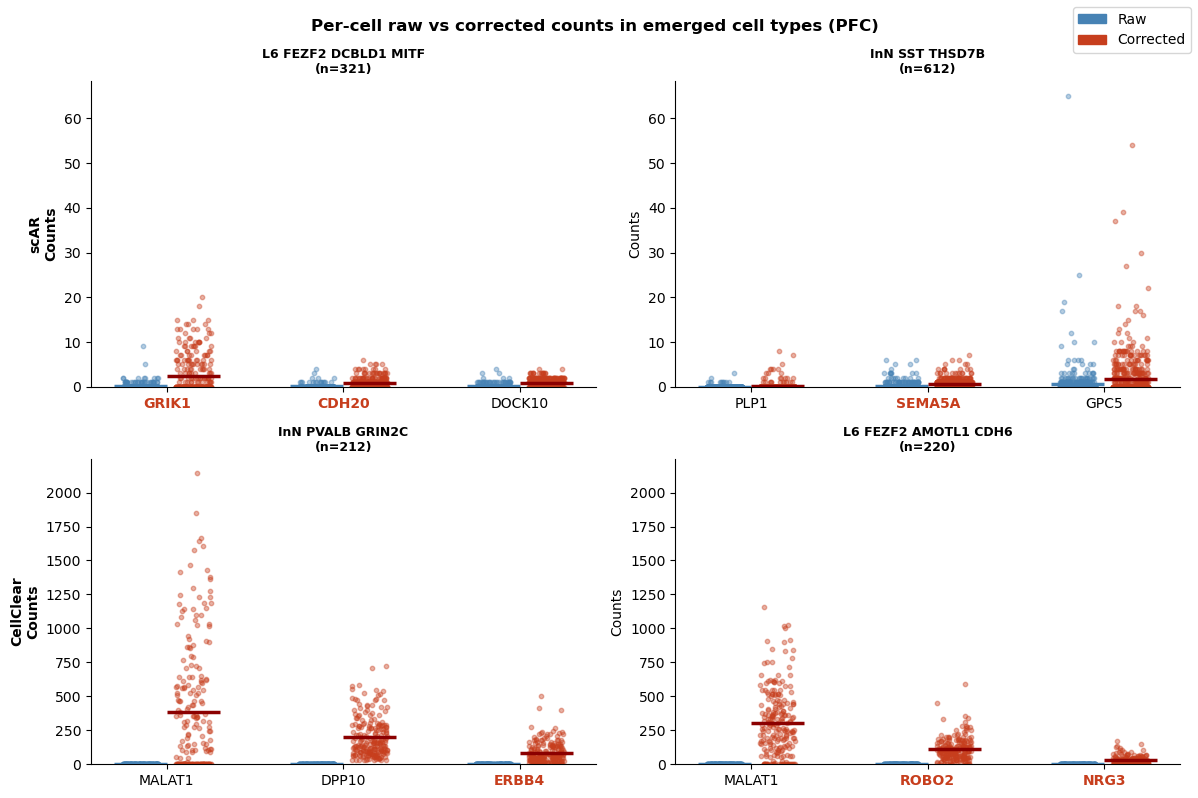

Supplementary Figure 4: Per-cell raw vs corrected counts in emerged PFC cell types

Strip plots showing per-cell raw (blue) and corrected (red) counts for selected genes in cell types that emerged after correction with scAR (top row) or CellClear (bottom row) in the prefrontal cortex snRNA-seq dataset but were absent from uncorrected data. Cell type labels and cell numbers per emerged type are indicated above each panel. scAR panels exemplify two emerged cell types: L6 FEZF2 DCBLD1 MITF (n=321; genes GRIK1, CDH20, DOCK10) and InN SST THSD7B (n=612; genes PLP1, SEMA5A, GPC5). CellClear panels exemplify InN PVALB GRIN2C (n=212; genes MALAT1, DPP10, ERBB4) and L6 FEZF2 AMOTL1 CDH6 (n=220; genes MALAT1, ROBO2, NRG3). Horizontal lines indicate per-gene means. Genes that are established markers of the respective cell type are visualized with bold red labels. Raw counts are near zero in the majority of affected cells, confirming that emerged cell types arise from artificial count inflation rather than denoising of low-but-present signal. Note the difference in y-axis scale between scAR (0–60 counts) and CellClear (0–2000 counts), reflecting the substantially larger magnitude of count inflation produced by CellClear.

### Supplementary Tables

###### Supplementary Table 1: Per-dataset ambient RNA removal metrics for all six tools across six hgmm species-mixing datasets.

Sensitivity: percentage of known inter-species ambient counts removed. Precision (extrapolated): accounts for unobservable intra-species ambient removal via symmetry-based extrapolation, capped at 100%. Specificity: computed from a per-species confusion matrix framework as TN / (TN + FP), where false positives are intra-species count removal exceeding the extrapolated ambient load. Total removed: percentage of all counts removed relative to uncorrected totals. Intra-species ratio: geometric mean of observed-to-expected intra-species removal ratios for human-in-human and mouse-in-mouse counts; a value of 1.0 indicates proportional removal relative to the extrapolated ambient load. The hgmm 1k dataset is included for completeness but was excluded from all aggregate metrics reported in Figure 2E due to markedly lower baseline inter-species contamination relative to all larger datasets, which is inconsistent with typical ambient RNA dynamics and would inflate apparent performance variability independent of tool behavior.

| Tool | Dataset | Sensitivity (%) | Precision, extrapolated (%) | Specificity (%) | Total removed (%) | Intra-species ratio  (geom. mean) |
| --- | --- | --- | --- | --- | --- | --- |
| CellBender | 10k | 97.3 | 96.4 | 99.9 | 2.1 | 1 |
| CellBender | 12k | 92.6 | 99.5 | 100 | 2.4 | 0.6 |
| CellBender | 1k | 93.3 | 56.5 | 99.7 | 0.7 | 2.5 |
| CellBender | 20k | 94.9 | 99.4 | 100 | 2.2 | 0.7 |
| CellBender | 5k | 94.9 | 100 | 100 | 1.5 | 0.5 |
| CellBender | 6k | 90.2 | 100 | 100 | 1.6 | 0.5 |
| CellClear | 10k | 85.8 | 5.4 | 60.2 | 41.1 | 36.6 |
| CellClear | 12k | 88.4 | 9 | 62.9 | 39.3 | 23.8 |
| CellClear | 1k | 85.2 | 1.2 | 62.4 | 37.9 | 136.7 |
| CellClear | 20k | 74.7 | 6.8 | 66.8 | 34.7 | 24.8 |
| CellClear | 5k | 90.1 | 6.8 | 67 | 34.5 | 33.3 |
| CellClear | 6k | 86.3 | 6.2 | 61 | 40.5 | 33.6 |
| DecontX | 10k | 62 | 100 | 100 | 1.4 | 0.6 |
| DecontX | 12k | 40.5 | 100 | 100 | 1.7 | 0.4 |
| DecontX | 1k | 68.9 | 74.9 | 99.9 | 0.4 | 0.7 |
| DecontX | 20k | 78.2 | 91.5 | 99.8 | 2.4 | 0.8 |
| DecontX | 5k | 42.6 | 100 | 100 | 1.2 | 0.4 |
| DecontX | 6k | 42.6 | 100 | 100 | 1.5 | 0.4 |
| SoupX | 10k | 50.4 | 100 | 100 | 1.2 | 0.6 |
| SoupX | 12k | 84.2 | 86.1 | 99.5 | 3.8 | 1.4 |
| SoupX | 1k | 89.1 | 39.3 | 99.4 | 1 | 3.9 |
| SoupX | 20k | 58 | 100 | 100 | 1.6 | 0.6 |
| SoupX | 5k | 83 | 95.7 | 99.9 | 2.3 | 1.2 |
| SoupX | 6k | 79.2 | 94.4 | 99.9 | 2.4 | 1.2 |
| scAR | 10k | 98.7 | 95.7 | 99.9 | 2.1 | 1 |
| scAR | 12k | 93.5 | 91.9 | 99.7 | 3.7 | 1.3 |
| scAR | 1k | 99.5 | 86.3 | 99.9 | 0.4 | 0.8 |
| scAR | 20k | 95.4 | 89.4 | 99.7 | 2.8 | 1.2 |
| scAR | 5k | 97.7 | 88.1 | 99.7 | 2.3 | 1.3 |
| scAR | 6k | 92.8 | 71.9 | 99 | 3.4 | 1.4 |
| scCDC | 10k | 16.5 | 87.2 | 99.9 | 1.1 | 0.9 |
| scCDC | 12k | 20.4 | 57.2 | 98.3 | 3.9 | 2.3 |
| scCDC | 1k | 6.2 | 11.6 | 98.3 | 1.9 | 7.1 |
| scCDC | 20k | 34.6 | 62.2 | 98.8 | 3 | 1.9 |
| scCDC | 5k | 23.6 | 43.5 | 98.2 | 3 | 2.7 |
| scCDC | 6k | 16.2 | 39.5 | 97.5 | 4 | 3.2 |

Supplementary Table 2: Pairwise hgmm sensitivity comparisons

Paired Wilcoxon signed-rank tests comparing sensitivity (ambient removed %) and total count removal (%) between CellBender, SoupX, DecontX, and scCDC across five hgmm datasets (1k excluded). Mean difference is tool 1 minus tool 2. Note: with n=5 paired observations, the minimum achievable p-value is 0.0313; single asterisk represents the maximum achievable significance level. scAR and CellClear were excluded from pairwise testing as they operate in distinct regimes (count inflation and near-complete count restructuring, respectively) that are addressed separately.

| Metric | Comparison | Mean difference (%) | U-statistic | p-value | Significance |
| --- | --- | --- | --- | --- | --- |
| Ambient removed | DecontX vs  SoupX | -18.17 | 4.0 | 0.2188 | ns |
| Ambient removed | DecontX vs CellBender | -38.07 | 0.0 | 0.0313 | * |
| Ambient removed | SoupX vs CellBender | -19.90 | 0.0 | 0.0313 | * |
| Ambient removed | DecontX vs  scCDC | 36.22 | 0.0 | 0. 0313 | * |
| Ambient removed | SoupX vs scCDC | 54.39 | 0.0 | 0.0313 | * |
| Ambient removed | CellBender vs  scCDC | 74.29 | 0.0 | 0.0313 | * |
| Total counts removed | DecontX vs  SoupX | -0.61 | 4.0 | 0.2185 | ns |
| Total counts removed | DecontX vs CellBender | -0.31 | 3.0 | 0.1563 | ns |
| Total counts removed | SoupX vs CellBender | 0.29 | 7.0 | 0.5625 | ns |
| Total counts removed | DecontX vs  scCDC | -1.37 | 1.0 | 0.0625 | ns |
| Total counts removed | SoupX vs scCDC | -0.77 | 2.0 | 0.0938 | ns |
| Total counts removed | CellBender vs  scCDC | -1.06 | 2.0 | 0.0938 | ns |

Supplementary Table 3: Marker gene enrichment Friedman test and post-hoc results

Friedman test statistics and post-hoc one-sided Wilcoxon signed-rank tests against zero (H₁: median log₂FC > 0) for each tool and dataset. BH correction applied across tools within each dataset. n=9 marker genes per dataset. scAR and CellClear excluded. DecontX° denotes DecontX run without the raw unfiltered count matrix.

| Dataset | Friedman χ² | Friedman p | Tool | Median log2FC | p (BH-corrected) | Significance |
| --- | --- | --- | --- | --- | --- | --- |
| PBMC | 18.467 | 0.0004 | CellBender | +0.332 | 0.0039 | ** |
| PBMC | 18.467 | 0.0004 | DecontX | +1.298 | 0.0078 | ** |
| PBMC | 18.467 | 0.0004 | SoupX | +0.217 | 0.0039 | ** |
| PBMC | 18.467 | 0.0004 | scCDC | +0.000 | 1.0000 | ns |
| WBC | 13.6 | 0.0087 | CellBender | +0.097 | 0.0033 | ** |
| WBC | 13.6 | 0.0087 | DecontX | +0.347 | 0.0033 | ** |
| WBC | 13.6 | 0.0087 | DecontX° | +0.191 | 0.0049 | ** |
| WBC | 13.6 | 0.0087 | SoupX | +0.065 | 0.0033 | ** |
| WBC | 13.6 | 0.0087 | scCDC | -0.319 | 0.9727 | ns |
| PFC | 23.133 | 0.0001 | CellBender | +0.528 | 0.0026 | ** |
| PFC | 23.133 | 0.0001 | DecontX | +0.976 | 0.0026 | ** |
| PFC | 23.133 | 0.0001 | SoupX | +0.114 | 0.0026 | ** |
| PFC | 23.133 | 0.0001 | scCDC | +0.000 | 1.0000 | ns |

Supplementary Table 4: Pairwise marker gene enrichment comparisons between tools

Pairwise two-sided Wilcoxon signed-rank tests on per-marker log₂ fold-change distributions across PBMC, WBC, and PFC datasets. n=9 marker genes per dataset. scAR and CellClear excluded due to count inflation invalidating ratio-based comparisons. No multiple testing correction applied across pairs; uncorrected p-values reported. DecontX° denotes DecontX run without the raw unfiltered count matrix.

| Dataset | Comparison | p-value (two-sided Wilcoxon) | Significance |
| --- | --- | --- | --- |
| PBMC | CellBender vs DecontX | 0.0742 | ns |
| PBMC | CellBender vs SoupX | 0.0078 | ** |
| PBMC | DecontX vs SoupX | 0.0195 | * |
| PBMC | CellBender vs scCDC | 0.0039 | ** |
| PBMC | DecontX vs scCDC | 0.0117 | * |
| PBMC | SoupX vs scCDC | 0.0039 | ** |
| WBC | CellBender vs DecontX | 0.0742 | ns |
| WBC | CellBender vs DecontX° | 0.4258 | ns |
| WBC | CellBender vs SoupX | 0.1289 | ns |
| WBC | DecontX vs DecontX° | 0.0742 | ns |
| WBC | DecontX vs SoupX | 0.0078 | ** |
| WBC | DecontX° vs SoupX | 0.0977 | ns |
| WBC | CellBender vs scCDC | 0.0547 | ns |
| WBC | DecontX vs scCDC | 0.0273 | * |
| WBC | DecontX° vs scCDC | 0.0391 | * |
| WBC | SoupX vs scCDC | 0.0742 | ns |
| PFC | CellBender vs DecontX | 0.0391 | * |
| PFC | CellBender vs SoupX | 0.0039 | ** |
| PFC | DecontX vs SoupX | 0.0078 | ** |
| PFC | CellBender vs scCDC | 0.0039 | ** |
| PFC | DecontX vs scCDC | 0.0039 | ** |
| PFC | SoupX vs scCDC | 0.0039 | ** |

Supplementary Table 5: Full CellTypist confidence score analysis

Per-tool CellTypist confidence score comparisons against uncorrected baseline across all datasets and annotation models. One-sided Mann-Whitney U tests (H₁: corrected > uncorrected) with Bonferroni correction across tools within each dataset-model combination; two-sided test applied where median difference was negative. All observed Cliff's δ values fall below the negligible threshold (|δ| < 0.147, Romano et al. 2006). Baseline uncorrected medians: PBMC coarse 1.0000, PBMC fine 0.9983, WBC coarse 0.9985, WBC fine 0.7074, PFC coarse 0.7581. scAR and CellClear excluded. DecontX° denotes DecontX run without the raw unfiltered count matrix.

| Data-set | Model | Tool | N cells | Median confidence | Δ Median | Cliff's δ | p (Bonfer-roni) | Significance |
| --- | --- | --- | --- | --- | --- | --- | --- | --- |
| PBMC | coarse | CellBender | 41428 | 0.999999 | 0.0000 | 0.144 | <0.0001 | ** |
| PBMC | coarse | DecontX | 38362 | 0.999999 | 0.0000 | 0.145 | <0.0001 | ** |
| PBMC | coarse | SoupX | 37560 | 0.999998 | 0.0000 | 0.073 | <0.0001 | ** |
| PBMC | coarse | scCDC | 38436 | 0.999992 | -0.0000 | -0.068 | <0.0001 | ** |
| PBMC | fine | CellBender | 41428 | 0.999430 | 0.0011 | 0.130 | <0.0001 | ** |
| PBMC | fine | DecontX | 38362 | 0.999064 | 0.0008 | 0.073 | <0.0001 | ** |
| PBMC | fine | SoupX | 37560 | 0.998975 | 0.0007 | 0.061 | <0.0001 | ** |
| PBMC | fine | scCDC | 38436 | 0.996750 | -0.0016 | -0.065 | <0.0001 | ** |
| WBC | coarse | CellBender | 118951 | 0.998792 | 0.0003 | 0.056 | <0.0001 | ** |
| WBC | coarse | DecontX | 121114 | 0.998734 | 0.0002 | 0.022 | <0.0001 | ** |
| WBC | coarse | DecontX° | 122541 | 0.998676 | 0.0002 | 0.014 | <0.0001 | ** |
| WBC | coarse | SoupX | 120632 | 0.998600 | 0.0001 | 0.009 | 0.0003 | ** |
| WBC | coarse | scCDC | 121872 | 0.998653 | 0.0001 | 0.020 | <0.0001 | ** |
| WBC | fine | CellBender | 118951 | 0.780884 | 0.0686 | 0.057 | <0.0001 | ** |
| WBC | fine | DecontX | 121114 | 0.705135 | -0.0071 | -0.008 | 0.0016 | ** |
| WBC | fine | DecontX° | 122541 | 0.715133 | 0.0029 | -0.001 | 1.0 | ns |
| WBC | fine | SoupX | 120632 | 0.716805 | 0.0045 | 0.004 | 0.3994 | ns |
| WBC | fine | scCDC | 121872 | 0.779606 | 0.0673 | 0.060 | <0.0001 | ** |
| PFC | coarse | CellBender | 152560 | 0.921215 | 0.1631 | 0.125 | <0.0001 | ** |
| PFC | coarse | DecontX | 161502 | 0.841427 | 0.0833 | 0.048 | <0.0001 | ** |
| PFC | coarse | SoupX | 158685 | 0.783475 | 0.0254 | 0.014 | <0.0001 | ** |
| PFC | coarse | scCDC | 158736 | 0.754777 | -0.0033 | 0.009 | <0.0001 | ** |
